# Acquisition of daptomycin resistance in patients results in decreased virulence in *Drosophila*

**DOI:** 10.1101/2024.10.07.616954

**Authors:** Brigitte Lamy, Frédéric Laurent, Carolina J. Simoes Da Silva, Ashima Wadhawan, Elizabeth V. K. Ledger, Camille Kolenda, Patricia Martins Simoes, Andrew M. Edwards, Marc S. Dionne

**Author notes:** Service de Microbiologie clinique,) APHP, Hôpitaux universitaires Paris Seine Saint-Denis, Bobigny, France UFR Santé, Médecine, Biologie humaine, Université Sorbonne Paris Nord (Paris 13, Bobigny, France. These authors also contributed equally to this work (Andrew M. Edwards, Marc S. Dionne).

## Abstract

*Staphylococcus aureus* can acquire antimicrobial resistance which in turn may affect its pathogenic potential. Using a panel of paired clinical isolates collected before and after daptomycin resistance acquisition, most frequently through single *mprF* mutation, we show a relationship between increasing daptomycin minimum inhibitory concentration and reduced virulence in a *Drosophila* systemic infection model. Analysing toxin production, *in vitro* bacterial growth characteristics, and cell surface properties, we failed to link daptomycin resistance-related attenuated virulence to either reduced virulence factor production, reduced fitness or to any of the cell surface characteristics investigated. Competition assays in *Drosophila* also did not support any altered ability in immune evasion. Instead, using a panel of mutant flies defective for various immune components, we show that this daptomycin resistance-related attenuated virulence is mostly explained by greater susceptibility to activity of *Drosophila* prophenoloxidase, a tyrosinase involved in melanization, but not to antimicrobial peptides or to Bomanin antimicrobial effectors. Further investigation could not link daptomycin resistance-related attenuation of virulence to a differential susceptibility to reactive oxygen species or to quinones prominently associated with phenoloxidase bacterial-killing activity. Taken together, it appears that daptomycin resistance attenuates *Staphylococcus aureus* virulence through an enhanced sensitivity to phenoloxidase based on a complex mechanism. Our study provides new insights in the understanding of the crosstalk between antimicrobial resistance, escape from immune killing, and virulence.

**Author summary:** Acquiring antimicrobial resistance can increase or decrease bacterial virulence. However, the mechanisms causing these resistance-virulence linked effects are unclear. Here, we bring new insights on the crosstalk between antimicrobial resistance and virulence. We characterized a panel of *Staphylococcus aureus* strains isolated from patients before and after resistance acquisition to the antibiotic daptomycin. Relative to the parental strain, resistant isolates most often varied by one single mutation, in a gene involved in the composition of the bacterial membrane, and these strains were much less virulent when fruit-flies were infected. Our results indicate that the difference of virulence is unrelated to changes in bacterial toxin production, bacterial growth, immune evasion or cell surface properties. Instead, resistant strains were more vulnerable to a host proenzyme involved in the antibacterial melanization response, an important response deployed throughout the arthropods. We predict that daptomycin resistance forces staphylococci to alter the composition of their cell surface. This alteration causes the bacteria to become more vulnerable to killing by melanization. Our results contribute to our understanding of the link between antimicrobial resistance and pathogenicity.

## Introduction

*Staphylococcus aureus* is a major cause of bacteremia. It has an estimated incidence of 9.3 to 65 cases/100,000 per year and a 30-day mortality rate between 18 and 29% [1,2]. Some sources of staphylococcal bacteremia stand out as difficult-to-treat infections (e.g., infective endocarditis or osteomyelitis), and the choice of treatment primarily relies on antimicrobial susceptibility [3–5]. While bacteremia caused by methicillin sensitive *S. aureus* (MSSA) is usually treated with antistaphylococcal penicillins, methicillin resistant *S. aureus* (MRSA) or by MSSA in penicillin-allergic patients [5,6] are treated with daptomycin, vancomycin, or linezolid [4,7,8]. However, treatment with both vancomycin and daptomycin can be complicated by the fact that resistance can arise via spontaneous mutations that occur during treatment [9,10].

Acquisition of antimicrobial resistance appears to be frequently associated with changes in virulence [11–13]. For example, vancomycin-intermediate *Staphylococcus aureus* can exhibit reduced fitness as a result of a thickened cell wall, enhanced capsular polysaccharide production, and dysfunction of the Agr virulence regulatory system [14–21]. However, the consequences of daptomycin resistance on fitness and virulence are less well established.

Daptomycin compromises bacterial surface integrity by targeting cell wall biosynthesis and membrane disruption, which may drive bacteria to alter cell surface characteristics and consequently affect immune host defence recognition [22,23]. Moreover, daptomycin shares a similar mechanism of action to host innate immune cationic antimicrobial peptides (AMP), which may facilitate cross-resistance between daptomycin and these important immune effectors [24–26]. Cross-resistance to host cationic AMPs, e.g., cathelicidin LL-37 or hNP-1, has been reported in some clinically derived daptomycin resistant *S. aureus* isolates in *in vitro* studies [24–26]. However, daptomycin resistance has been reported to result in attenuated virulence in a zebrafish model (e.g., [26]).

*Drosophila melanogaster* is a well-established model for the study of antibacterial immunity [27]. *Staphylococcus aureus* peptidoglycan is detected via the circulating pattern recognition receptor PGRP-SA, resulting in activation of the Toll signalling pathway as well as an antibacterial melanization response [28]. The Toll pathway drives production of antimicrobial peptides, while the melanization response depends on phenoloxidase activity. Of these responses, melanization appears to be the most important mechanism of defense against *Staphylococcus aureus* infection [29]. This response depends on conversion of prophenoloxidase to active phenoloxidase by serine protease cleavage. Phenoloxidase then converts tyrosine to dopaquinone, which polymerizes to form melanin. It is unclear whether dopaquinone, associated reactive oxygen species, or melanin itself is the relevant killing mechanism [30,31].

It is commonly acknowledged that some host immune effectors and antimicrobials share common mechanisms of action, suggesting potential mechanistic links between virulence and antimicrobial resistance [32,33]. Although there is growing evidence of a trade-off between daptomycin resistance and pathogenic potential, the associated mechanisms are still poorly understood. This is further confounded by the fact that investigation is often limited to single clinical paired strains or to laboratory generated daptomycin-resistant strains [21,26,34,35]. Thus, the ability to generalise previous findings is limited by the use of one or very few strain pairs. Moreover, many studies focus on the correlation between AMR and *in vitro* proxies for virulence, such as gene expression, but the impact of these changes on *in vivo* virulence is often unexplored. Finally, the molecular determinants underlying the interaction between antimicrobial resistance and pathogenic potential are often unclear. Understanding these determinants is informative regarding the fundamental biology of treatment and pathogenesis and may help identify new antimicrobial strategies to either restore antimicrobial sensitivity or reduce virulence.

Here, we were interested in exploring the trade-off between daptomycin resistance and staphylococcal virulence as a strategy to deepen our understanding of staphylococcal pathogenicity, and thus inform efforts to devise new antimicrobial strategies. Using a panel of *S. aureus* clinical strains recovered from bloodstream infection before and after daptomycin resistance acquisition, we investigated the impact of daptomycin resistance on virulence using a *Drosophila* systemic infection model to decipher crosstalk between daptomycin resistance and staphylococcal pathogenesis. We also investigated the impact of daptomycin resistance on interaction between *S. aureus* and innate immunity. This led to the discovery that daptomycin resistance is associated with increased susceptibility to phenoloxidase-mediated killing.

## Results

### The acquisition of daptomycin resistance during treatment results in reduced virulence

To understand the consequences of *in vivo* acquired daptomycin resistance on staphylococcal virulence, we collected 11 pairs of isolates from patients with *S. aureus* endocarditis in whom daptomycin resistance arose during treatment. Isolates collected before treatment were all susceptible to daptomycin and those collected after were resistant to the lipopeptide antibiotic.

In keeping with the literature, daptomycin resistance was associated with 2 to 8-fold increases in daptomycin MIC (from 0.125-0.5 to 1-2 mg/L) and oxacillin MIC decreases in 9/11 pairs (2 to 64-fold reduction), whilst vancomycin MICs were unchanged (**Table 1**) [36–39]. Whole genome sequencing found 7/11 pairs in which susceptible and resistant strains differed only by single mutations in the multipeptide resistance factor (*mprF)* gene commonly associated with daptomycin resistance [40,41]. One pair (pair D) had acquired mutations in *mprF* and *rpoB*, whilst pair J had only acquired a mutation in *rpoB*. Pairs B and H had acquired multiple insertions, deletions and single nucleotide polymorphism in addition to *mprF* mutations (n=3 and n=13, respectively) (**Table 1**). Thus, the isolates assembled here are representative of previously reported clinical daptomycin resistant strains [10,42,43]. Furthermore, since many isolates differed by a single mutation, we were able to investigate the impact of daptomycin resistance on virulence in the absence of potentially confounding mutations.

**Table 1.**
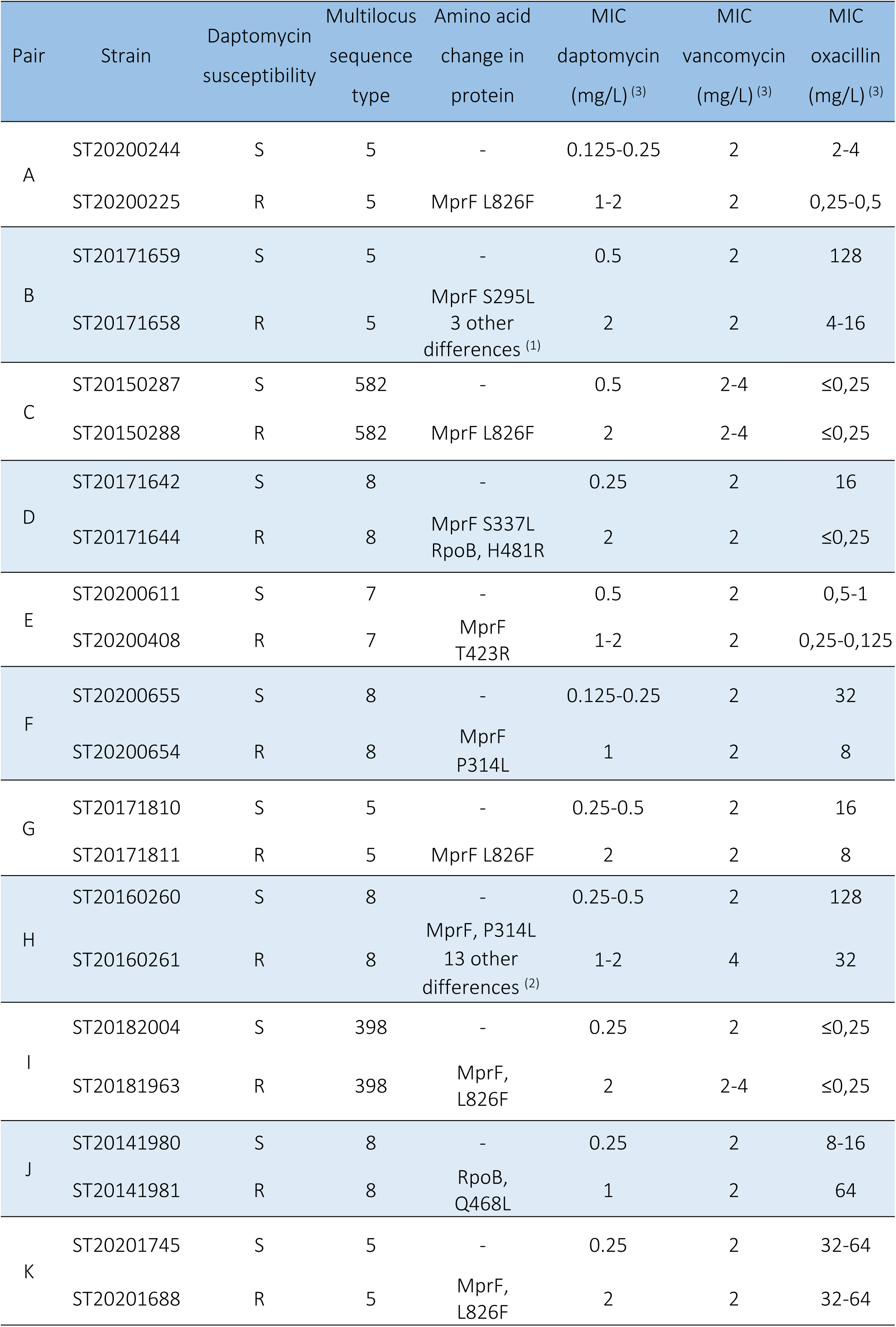
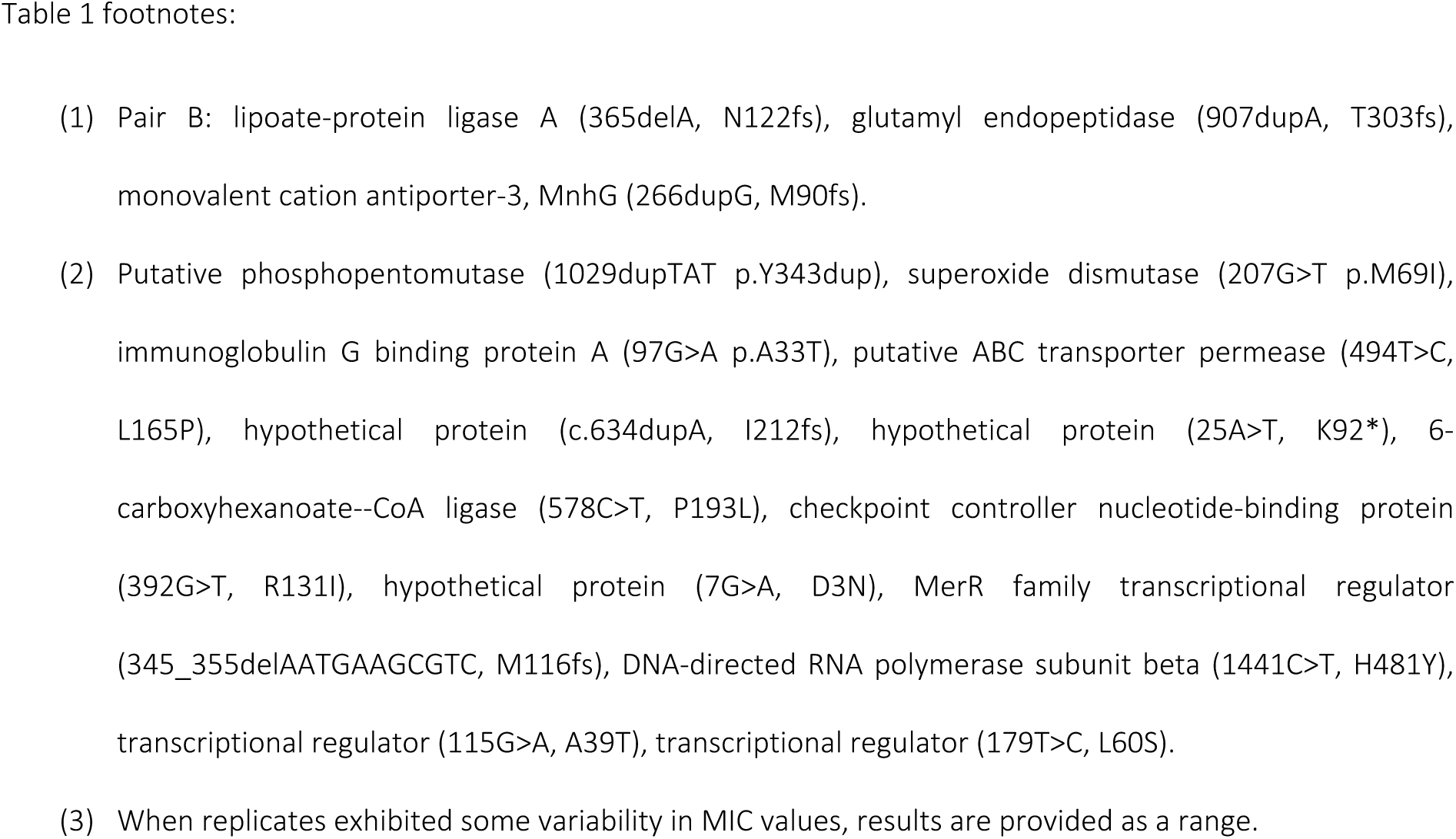
Characteristics of the clinical strains included in this study.

| Pair | Strain | Daptomycin<br>susceptibility | Multilocus<br>sequence<br>type | Amino acid<br>change in<br>protein | MIC<br>daptomycin<br>(mg/L) <sup>(3)</sup> | MIC<br>vancomycin<br>(mg/L) <sup>(3)</sup> | MIC<br>oxacillin<br>(mg/L) <sup>(3)</sup> |
| --- | --- | --- | --- | --- | --- | --- | --- |
| A | ST20200244 | S | 5 | - | 0.125-0.25 | 2 | 2-4 |
|  | ST20200225 | R | 5 | MprF L826F | 1-2 | 2 | 0,25-0,5 |
| B | ST20171659 | S | 5 | - | 0.5 | 2 | 128 |
|  | ST20171658 | R | 5 | MprF S295L<br>3 other<br>differences <sup>(1)</sup> | 2 | 2 | 4-16 |
| C | ST20150287 | S | 582 | - | 0.5 | 2-4 | ≤0,25 |
|  | ST20150288 | R | 582 | MprF L826F | 2 | 2-4 | ≤0,25 |
| D | ST20171642 | S | 8 | - | 0.25 | 2 | 16 |
|  | ST20171644 | R | 8 | MprF S337L<br>RpoB, H481R | 2 | 2 | ≤0,25 |
| E | ST20200611 | S | 7 | - | 0.5 | 2 | 0,5-1 |
|  | ST20200408 | R | 7 | MprF<br>T423R | 1-2 | 2 | 0,25-0,125 |
| F | ST20200655 | S | 8 | - | 0.125-0.25 | 2 | 32 |
|  | ST20200654 | R | 8 | MprF<br>P314L | 1 | 2 | 8 |
| G | ST20171810 | S | 5 | - | 0.25-0.5 | 2 | 16 |
|  | ST20171811 | R | 5 | MprF L826F | 2 | 2 | 8 |
| H | ST20160260 | S | 8 | - | 0.25-0.5 | 2 | 128 |
|  | ST20160261 | R | 8 | MprF, P314L<br>13 other<br>differences <sup>(2)</sup> | 1-2 | 4 | 32 |
| I | ST20182004 | S | 398 | - | 0.25 | 2 | ≤0,25 |
|  | ST20181963 | R | 398 | MprF,<br>L826F | 2 | 2-4 | ≤0,25 |
| J | ST20141980 | S | 8 | - | 0.25 | 2 | 8-16 |
|  | ST20141981 | R | 8 | RpoB,<br>Q468L | 1 | 2 | 64 |
| K | ST20201745 | S | 5 | - | 0.25 | 2 | 32-64 |
|  | ST20201688 | R | 5 | MprF,<br>L826F | 2 | 2 | 32-64 |

- (1) Pair B: lipote-protein ligase A (365delA, N122fs), glutamyl endopeptidase (907dupA, T303fs), monovalent cation antiporter-3, MnhG (266dupG, M90fs). - (2) Putative phosphopentomutase (1029dupTAT p.Y343dup), superoxide dismutase (207G>T p.M69I), immunoglobulin G binding protein A (97G>A p.A33T), putative ABC transporter permease (494T>C, L165P), hypothetical protein (c.634dupA, I212fs), hypothetical protein (25A>T, K92\*), 6-carboxyhexanoate--CoA ligase (578C>T, P193L), checkpoint controller nucleotide-binding protein (392G>T, R131I), hypothetical protein (7G>A, D3N), MerR family transcriptional regulator (345\_355delAATGAAGCGTC, M116fs), DNA-directed RNA polymerase subunit beta (1441C>T, H481Y), transcriptional regulator (115G>A, A39T), transcriptional regulator (179T>C, L60S). - (3) When replicates exhibited some variability in MIC values, results are provided as a range.

Therefore, we proceeded to test the virulence of our strains in *Drosophila*. Most (8/11) resistant strains showed a strong reduction in host killing in comparison with their paired susceptible strain (**Fig 1A**). Only pairs J and K exhibited no difference in virulence between resistant and paired susceptible strains (**Fig 1A**). The lack of difference in virulence between the strains in pair K was surprising since pairs A, C, G, I and K differ by the same L826F amino acid substitution in MprF (Table 1). The different consequences observed in strains carrying the same mutation may be a result of the differences in genetic background among the strains.

**Fig 1.**
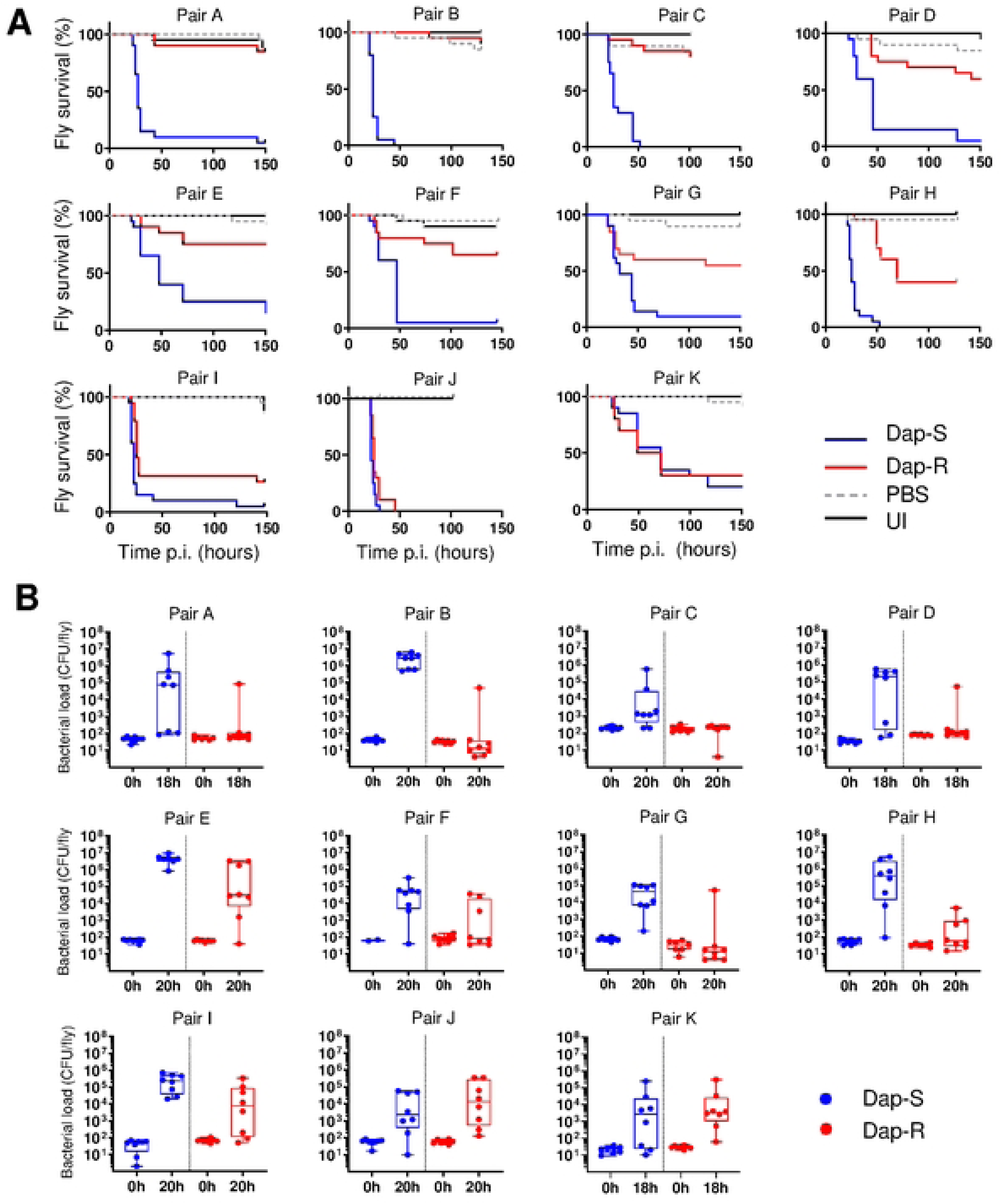
Daptomycin resistant strains exhibit lower virulence and reduced growth in the fly relative to susceptible isolates. (A) Survival of wild-type (*w^1118^*) flies infected with Dap-S strain and Dap-R paired isolate. Each graph corresponds to one experiment with 20 flies per condition and is representative of at least two independent experiments. P-values were calculated using the Log-rank test. P-values for survival differences with Dap-R isolate vs Dap-S paired isolate: Pairs A-F and H, p≤0.0001, pair G, p=0.007, pair I, p=0.0006, pair J, p=0.046, pair K, p=0.636; PBS and UI, p>0.05. (B) Strain colony counts at 0h (input inoculum) and 18 to 20h after injection of wild type (*w^1118^*) flies with Dap-S strain and Dap-R paired isolate. Bacterial counts presented correspond to one experiment and are representative of two independent experiments. P-values for bacterial numbers at 18-20 h post infection (Dap-R vs Dap-S paired strain) were calculated using the Mann-Whitney test: pair A, p=0.0104; pair B, p=0.0002; pair C, p=0.0342; pair D, p=0.0779; pair E, p=0.004; pair F, p=0.0277; pair G, p=0.0017; pair H, p=0.0011; pair I, p=0.0207; pair J, p=0.36; pair K, p=0.64. Dap-S, daptomycin susceptible; Dap-R, Daptomycin resistant ; UI, uninjected flies.

The observation that all but three of the daptomycin resistant strains exhibited a clear attenuation of virulence relative to their matched susceptible isolate suggests a fundamental link between daptomycin resistance and *Staphylococcus aureus* pathogenesis.

We next determined whether these differences in virulence reflected differences in the ability of the bacteria to grow in the host. This was assayed by quantifying viable bacterial counts in infected animals. We found that increased host mortality correlated with elevated viable bacterial numbers 18 to 20 hours after infection (**Fig 1B**). In most cases, the fly was able to control growth of daptomycin resistant strains but was unable to control the paired daptomycin susceptible strain. This was not a general proliferation defect since the paired resistant and susceptible strains did not differ in growth rates in four different laboratory media (**S1 Fig**).

Daptomycin resistance is frequently associated with changes in various cell surface characteristics[44]. We examined several cell envelope properties of the isolates, including membrane fluidity, cell surface charge, and cell wall thickness. We also examined production of the carotenoid pigment staphyloxanthin, as this has been shown to confer protection against host defences such as reactive oxygen species. However, there were no significant differences between daptomycin susceptible and resistant strains for any of these properties (**Fig 2 and** Figures **S2-S5**).

**Fig 2.**
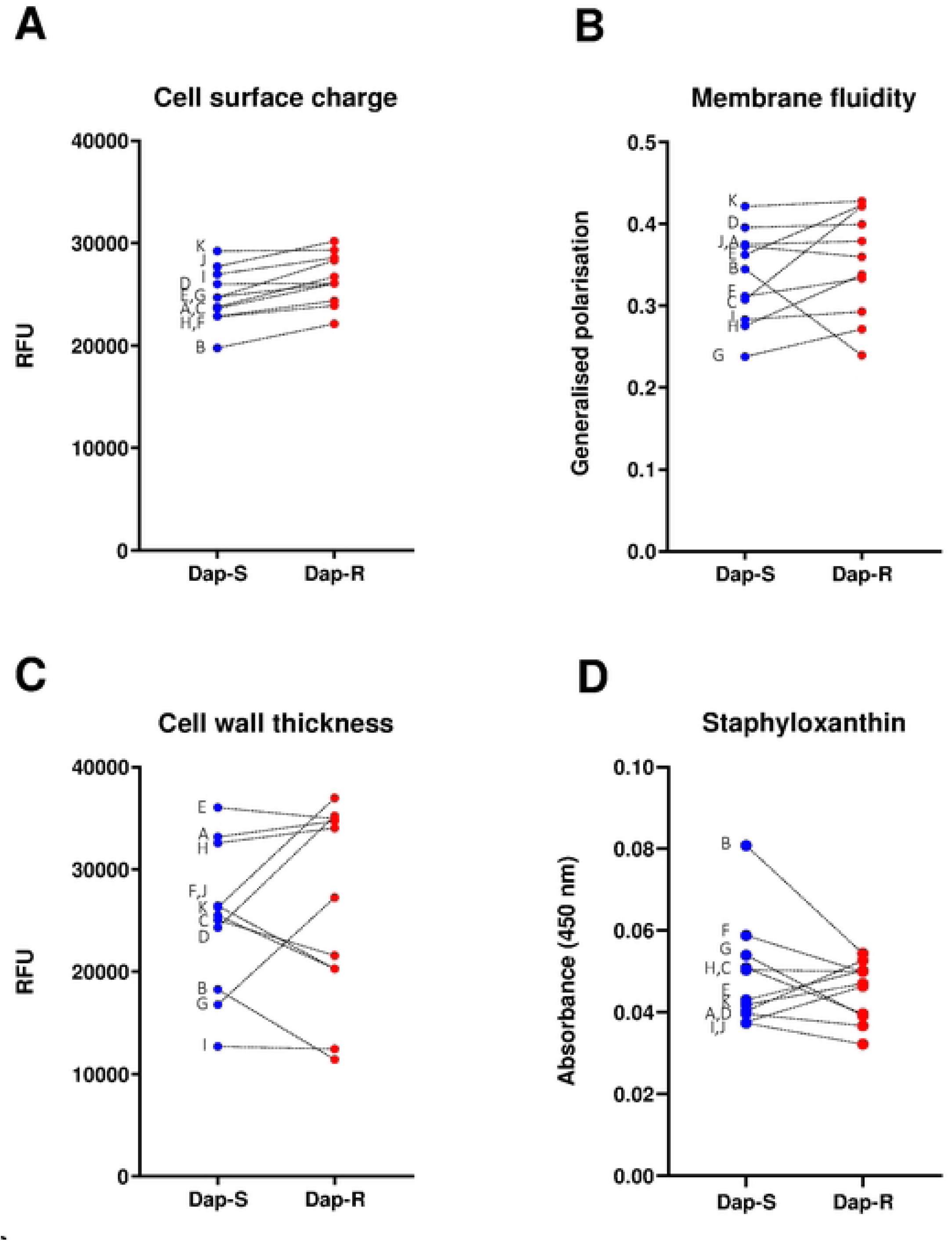
Cell surface properties do not correlate with daptomycin resistance or virulence. Data from three independent experiments. Dashed lines link paired strains; letters indicate pair name. Plots represent mean, error bars are omitted for clarity. (A) Cell surface charge, p=0.004, see also S2 Fig for detailed results; (B) fluidity of cell membrane, p>0.05, see also S3 Fig for detailed results, (C), cell wall thickness, p>0.05, see also S4 Fig for detailed results; (D) cell surface carotenoid content, p>0.05, see also S5 Fig for detailed results. Data were analysed by a two-tailed paired Student’s t-test. Dap-S, daptomycin susceptible; Dap-R, daptomycin resistant. RFU, relative fluorescence units.

### Reduced virulence of daptomycin resistant isolates is not due to altered virulence factor production

*S. aureus* produces a wide range of different virulence factors, including several cytolytic toxins expressed under the control of the Agr quorum sensing system [45]. Since toxin production is important for both human and *Drosophila* infection [45–48], we hypothesised that the lack of virulence of daptomycin resistant isolates could be due to reduced virulence factor production.

To test this, we assessed the production of exotoxins by quantifying haemolytic activity as this is a marker of activity of the Agr quorum-sensing system [20]. Most strains exhibited haemolytic activity after 4h growth in broth, consistent with secretion of haemolytic toxins resulting from Agr activity. While haemolytic activity varied among pairs, we did not detect a consistent difference between individual isolates within pairs (**Fig 3A**). We then extended this analysis to proteins secreted by the bacterial cells (the exoproteome), focusing on pairs that carried the same MprF amino acid substitution (L826F, pairs A, C, G, I, K). Again, we found variation in SDS-PAGE profile among pairs, but no consistent difference between individual isolates within pairs (**Fig 3B**). Collectively, these data suggest that reduced virulence exhibited by daptomycin resistant strains is not related to differences in exotoxin production.

**Fig 3.**
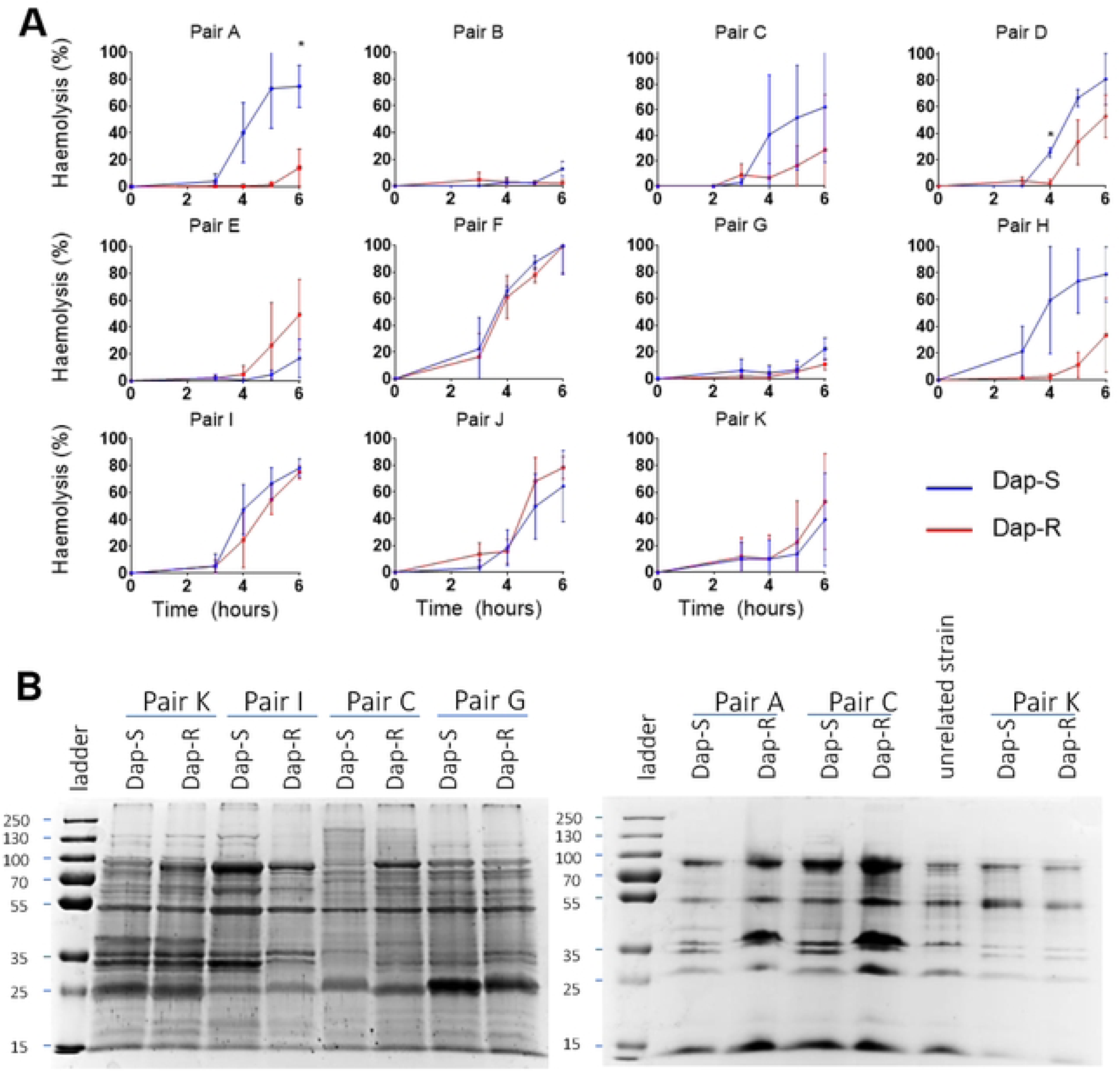
Virulence changes are not driven by changes in toxin secretion. (A) Haemolytic activity of culture supernatants from all paired strains. Data were analysed by a two-way ANOVA with Sidak’s post hoc tests. P-values for difference in haemolysis activity produced by Dap-S vs Dap-R. P-values <0.05 are indicated with *. Pair A, p=0.0375 (6 h), Pair D, p=0.0058 (4 h), all other P-values >0.05. (B) 10% SDS-PAGE of 10-fold concentrated supernatants from 4h cultures. All pairs shown on this gel exhibit the same MprF amino acid substitution (L826F). Gel representative of two independent assays. Dap-S, daptomycin susceptible; Dap-R, Daptomycin resistant.

### Reduced virulence of daptomycin resistant isolates is not due to increased immune activation

In *Drosophila* the Toll signalling pathway and the antibacterial melanization response rely on a common peptidoglycan sensing mechanism [29]. The melanization response is the primary immune effector mechanism functional against *S. aureus*, but the production of antimicrobial peptides driven by Toll pathway activation also contributes to host defence against this bacterium, and the expression of peptide genes is a well-established marker of Toll activation [29].

To test whether daptomycin susceptible and resistant strains drove different levels of immune response activation, we measured expression of genes expressed in response to immune activation for strain pairs A and D. As expected, PBS injection induced *Drosomycin* (*Drs*), *Metchnikowin* (*Mtk*), *Attacin A* (*AttA*), and *Bomanin S2* (*BomS2*) expression (**Fig 4**). Daptomycin susceptible and resistant isolates induced the expression of these peptides at levels similar to or higher than PBS injection. There was no strong difference between daptomycin susceptible and resistant isolates in the expression of any peptide genes tested. These results support the idea that the difference in virulence between paired isolates is not derived from differential ability to evade detection by the mechanisms upstream of Toll activation and melanization.

**Fig 4.**
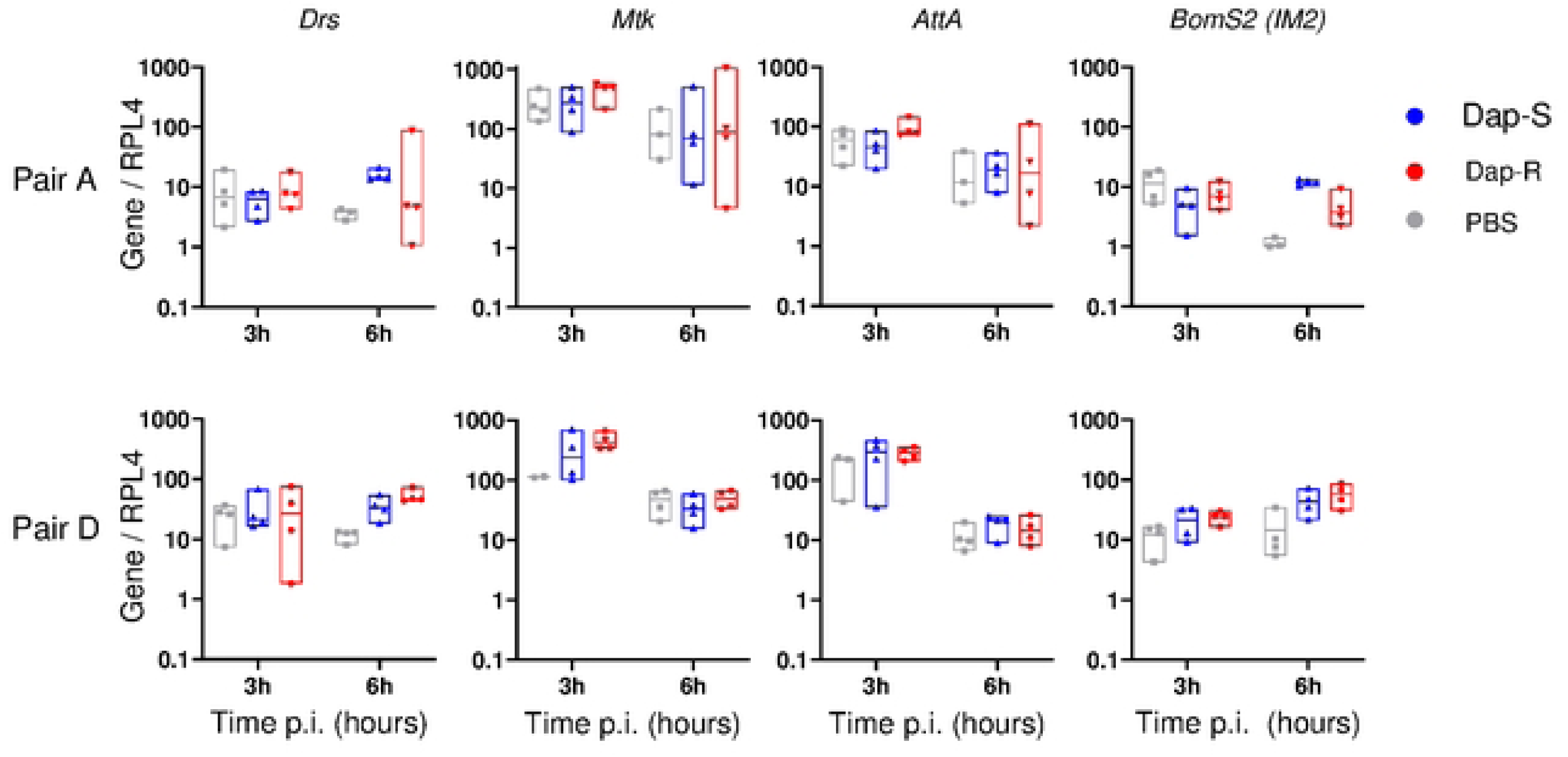
AMP genes are expressed at similar levels following infection with daptomycin resistant and susceptible strains. Host AMP gene expression 3h and 6h after fly infection. Assessed AMPs are representative of immune pathways activated by bacterial infection. Expression levels of AMPs are shown normalized to *Rpl4* and then to the mean value for uninjected flies. Data were analysed by ANOVA followed by Tukey’s post hoc test for multiple comparisons. *Drs*, pair A, 3h: p>0.05 for all comparisons; 6h: p>0.05 for all comparisons; pair D, 3h: p>0.05 for all comparisons; 6h: Dap-S and PBS, p=0.0523; Dap-R and PBS, p=0.024; Dap-S and Dap-R, p>0.05. *Mtk* and *AttA*, 3h and 6h: p>0.05 for all comparisons (all pairs). *BomS2*: pair A, 3h: p>0.05 for all comparisons; 6h: Dap-S and PBS, p=0.0004; Dap-R and PBS, p>0.05; Dap-S and Dap-R, p=0.0032; pair D, 3h: p>0.05 for all comparisons; 6h: Dap-S and PBS, p>0.05; Dap-R and PBS, p=0.0341; Dap-S and Dap-R, p>0.05. Dap-S, daptomycin susceptible; Dap-R, daptomycin resistant. Drs, Drosomycin; Att: attacin A; Mtk: Metchnikowin, BomS2: Bomanin S2.

It remained possible that the daptomycin susceptible strains could interfere with some aspect of the immune response other than antimicrobial peptide transcription. In this case, co-infection with daptomycin susceptible and resistant strains would allow daptomycin resistant strains to grow *in vivo* to a similar degree to their paired susceptible strain. Therefore, we co-infected *Drosophila* with 1:1 mixtures of susceptible and resistant bacteria and we measured the numbers of viable susceptible and resistant bacteria 20h after infection. This was explored for pairs A and D.

As before, in single strain infections, daptomycin susceptible strains grew *in vivo*, whilst daptomycin resistant bacteria did not. In co-infection experiments, the presence of daptomycin susceptible bacteria was not able to rescue the ability of daptomycin resistant bacteria to grow *in vivo* (**Figs 5A and 5B**).

**Fig 5.**
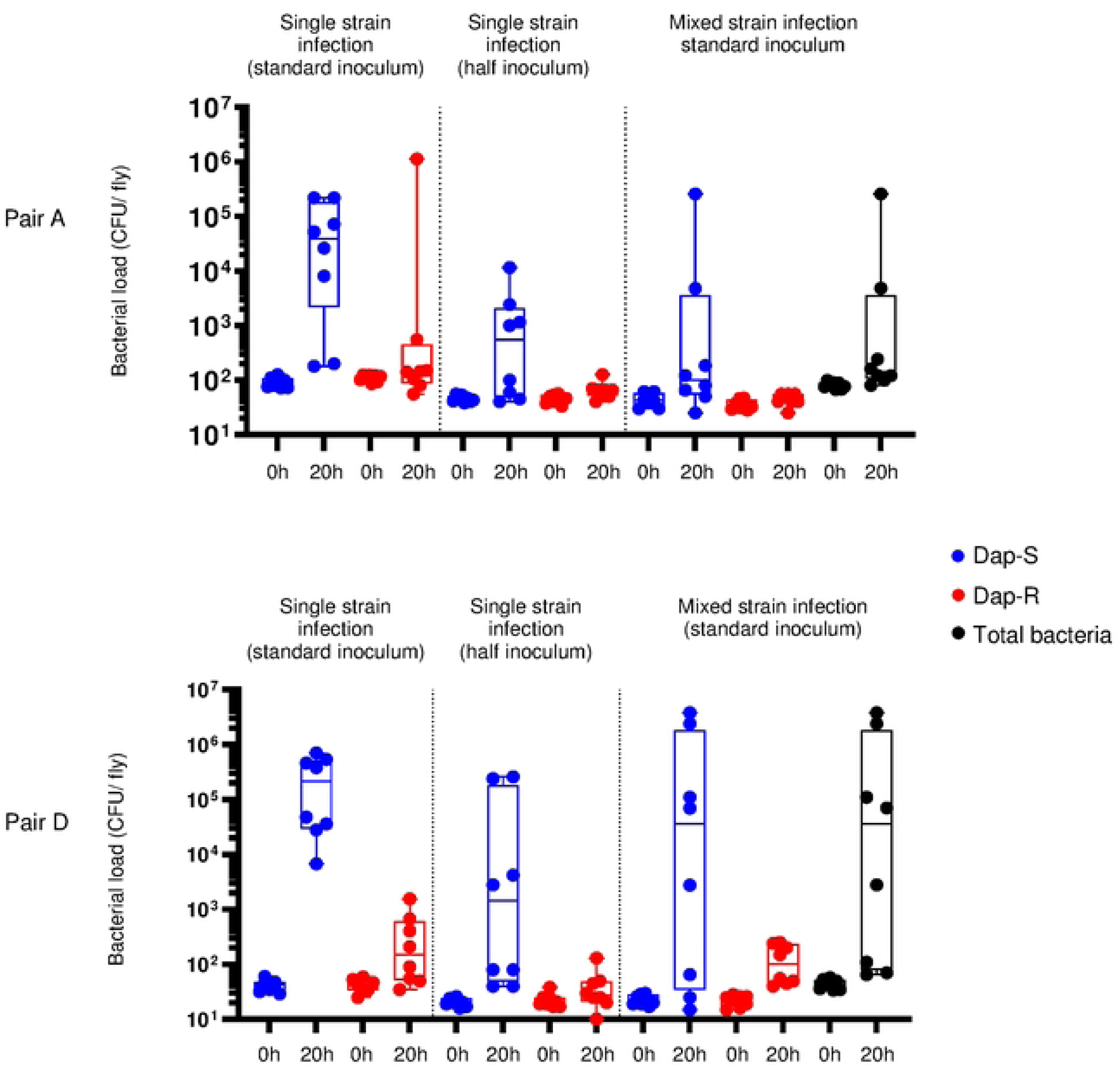
Daptomycin susceptible strains do not confer protection on co-infected daptomycin resistant bacteria. Strain colony counts at 0 h (input inoculum) and 20 h after injection of wild type (*w^1118^*) flies with Dap-S strain, Dap-R paired strain or Dap-S strain mixed with Dap-R paired strain. The ratio Dap-S:Dap-R in the mixed condition is 1:1; the total inoculum in co-infection experiments is the same as the standard inoculum used in other experiments. Bacterial counts presented correspond to one experiment representative of two independent experiments. P-values for bacterial numbers at 20 h post infection (Dap-R vs Dap-S paired strain) were calculated using a Kruskall-Wallis test with Dunn’s post hoc test for multiple comparisons. (A) Pair A, Dap-S/Dap-R co-infection assay. Single strain infections were performed with an average input inoculum of 92 CFU/fly (Dap-S) and 75 CFU/fly (Dap-R, standard inoculum), and of 26 CFU/fly (Dap-R) and 33 CFU/fly (Dap-S, half inoculum), mixed strain infections were performed with an average of 36 CFU/fly (Dap-S) and 35 CFU/fly (Dap-R), which resulted in an average total inoculum of 71 CFU/fly. At 20h, all P-values>0.05, except: Dap-S vs Dap-R, standard inoculum: p=0.05; Dap-S, half inoculum vs Dap-R, co-infection: p=0.045. (B) Pair D, Dap-S/Dap-R co-infection assay. Single strain infections were performed with an average input inoculum of 42 CFU/fly (Dap-S) and 37 CFU/fly (Dap-R, standard inoculum), and of 20 CFU/fly (Dap-S) and 22 CFU/fly (Dap-R, half inoculum), mixed strain infections were performed with an average of 23 CFU/fly (Dap-S) and 22 CFU/fly (Dap-R), resulting in an average total inoculum of 45 CFU/fly. At 20h, all P-values>0.05, except: Dap-S vs Dap-R, standard inoculum: p=0.004; Dap-S vs Dap-R, half inoculum: p=0.053; Dap-S vs Dap-R, co-infection: p=0.046.

Taken together, these data indicated that the reduced virulence of daptomycin resistant isolates relative to their matched daptomycin susceptible partner was not due to enhanced immune activation.

### Daptomycin resistant strains are more susceptible to killing by melanization but not to antimicrobial peptides

Since daptomycin susceptible strains were unable to protect resistant strains from host immune defences, we assumed that susceptible strains were more resistant to host immune killing mechanisms than resistant isolates (**Fig 4 and Fig 5**). To test this, we first investigated whether Toll-dependent antimicrobial peptides could be responsible for this difference by measuring survival after infection of *Dif*-mutant flies, which lack almost all Toll-dependent inducible gene expression, and Bomanin-deficient flies, which lack a family of peptides important in defence against some Gram-positive bacteria [49,50]. However, the absence of *Dif* or of Bomanins did not affect the survival difference between paired susceptible and resistant strains (**Fig 6A** and **S6 Fig**). We then infected transgenic flies expressing *Toll^10B^*, a constitutive allele of *Toll* [51]. As had been seen in wild-type flies, daptomycin susceptible strains rapidly killed flies with constitutively active Toll signalling, while their resistant counterparts were significantly attenuated (**Fig 6B**). Taken together, these results indicate that that daptomycin resistance -related attenuated virulence is not associated with increased sensitivity to Toll-dependent antimicrobial peptides or Bomanins.

**Fig 6.**
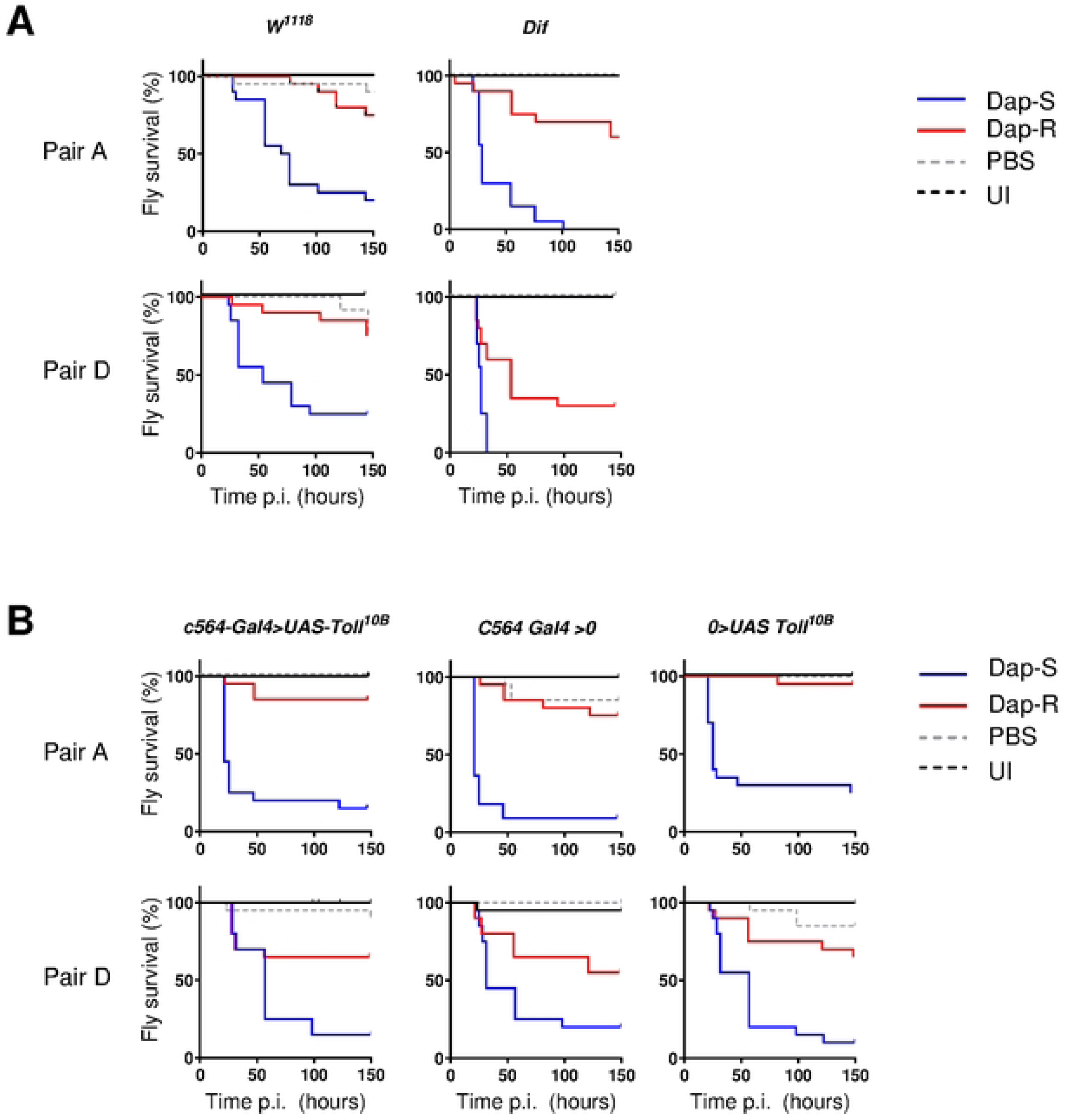
Daptomycin resistance does not increase sensitivity to Toll-dependent antimicrobial effectors. Data presented for two pairs, pair A and pair D. (A) Survival of wild type (*w^1118^*) and *Dif*-mutant flies infected with Dap-S or Dap-R paired strains. Survival curves correspond to one assay representative of two independent experiments. (B) Survival of control (*C564-Gal4>0* and *0>UAS-Toll^10B^*) and *Toll^10b^*-expressing (*C564-Gal4>UAS-Toll^10B^*) flies infected with Dap-S and Dap-R paired strains. Survival curves correspond to one assay representative of two independent experiments.

We next investigated whether differential sensitivity to melanization was responsible for the virulence difference between daptomycin susceptible and resistant strains. We tested this by determining the sensitivity of flies lacking the prophenoloxidases *PPO1* and *PPO2*; these enzymes are required for immune-induced melanization and are important for defense against *S. aureus* [52]. We observed that the difference in survival between flies infected with daptomycin susceptible and resistant paired strains was consistently reduced in *PPO1^Δ^*, *PPO2^Δ^* double mutants than in wild-type controls (**Fig 7A**, **S7A Fig**). This effect was also seen in bacterial loads: daptomycin susceptible strains grew to equal numbers in wild-type and *PPO1^Δ^, PPO2^Δ^* flies, while daptomycin resistant strains exhibited little growth in wild-type flies but grew in *PPO1^Δ^, PPO2^Δ^* flies to similar numbers to susceptible strains (**Fig 7B**). These findings demonstrate that the daptomycin resistance -related loss of virulence results from differential sensitivity to killing by phenoloxidase.

**Fig 7.**
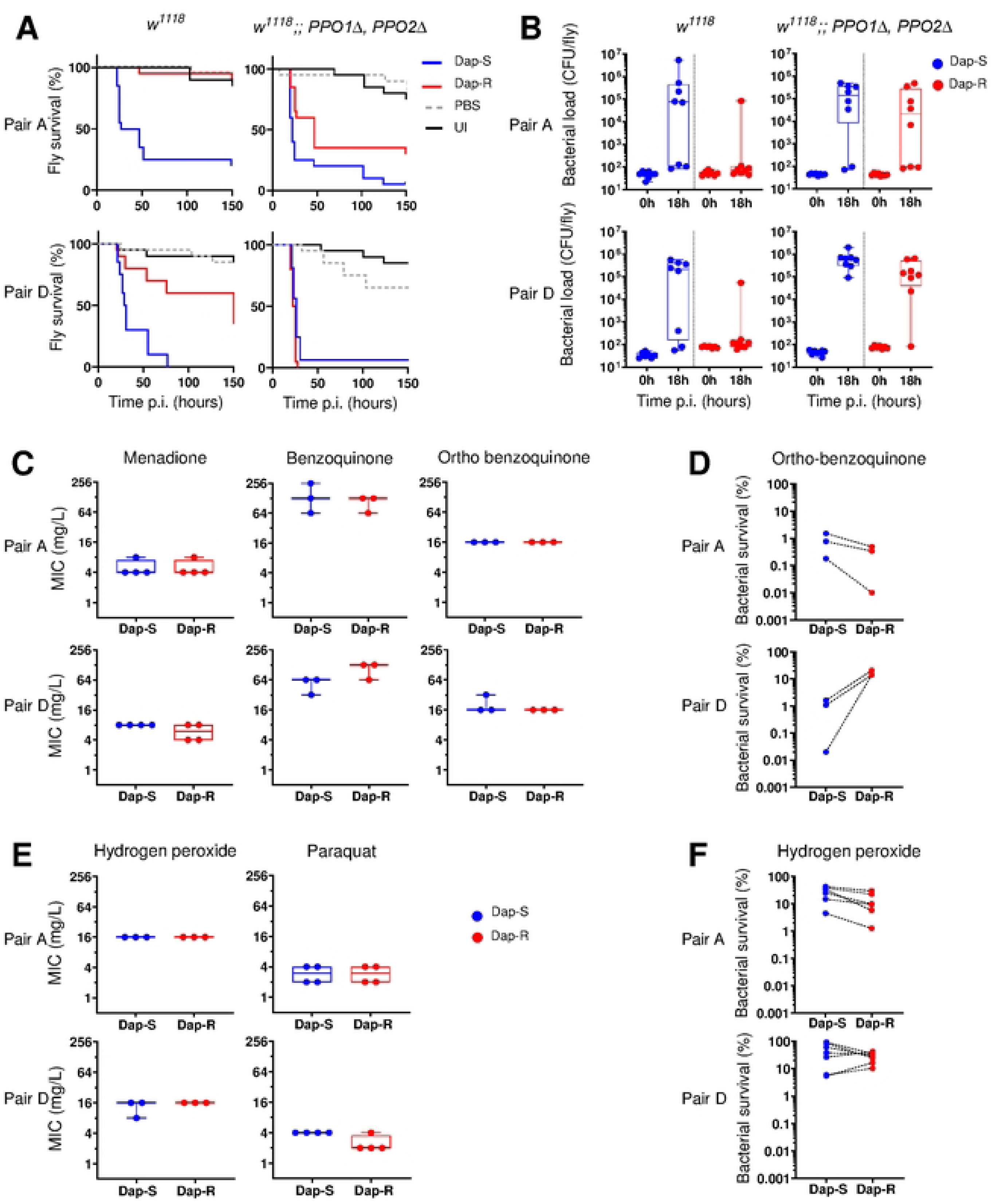
Dap-R staphylococcal virulence is affected by prophenoloxidase but cells are not more susceptible to quinones and reactive oxygen species. Data presented for 2 pairs, pair A and pair D. (A) Survival of wild type (*w^1118^*) and *w^1118^; PPO1^Δ^ PPO2^Δ^* flies infected with either Dap-S strain or Dap-R paired strain. Survival analysis reveals similar mortality of *PPO1^Δ^ PPO2^Δ^*double mutant flies whether infected with Dap-S or Dap-R strains. Survival curves correspond to one assay representative of at least two independent experiments. See also supplementary figure S7A for other pairs. (B) Bacterial numbers at 0 h (input inoculum) and 18 h after injection of wild type (*w^1118^*) or *w^1118^; PPO1^Δ^ PPO2^Δ^* flies with Dap-S strain and Dap-R paired strain. Counts from one assay representative of at least two independent experiments. Bacterial load is similar Dap-R and Dap-S isolates in *PPO1^Δ^ PPO2^Δ^* flies (pair A, median bacterial numbers at 18h : in *w^1118^*, Dap-S, 7.6x10^5^ CFU/fly vs Dap-R, 72 CFU/fly; in *PPO1^Δ^ PPO2^Δ^,* Dap-S, 1.4 x10^5^ CFU/fly vs Dap-R, 2.1 x10^4^ CFU/fly; Mann-Whitney test, p=0.0104 (*w^1118^*), p=0.4609 (*PPO1^Δ^ PPO2^Δ^*); pair D, median bacterial numbers at 18h in *w^1118^*: Dap-S, 2.1 x10^5^ CFU/fly vs Dap-R, 112 CFU/fly; in *PPO1^Δ^ PPO2^Δ^:* Dap-S, 6.0 x10^5^ CFU/fly vs Dap-R, 1.3 x10^5^ CFU/fly; Mann-Whitney test, p=0.0779 (*w^1118^*), p=0.0379 (*PPO1^Δ^ PPO2^Δ^*). CFU, colony forming unit. (C) Minimum inhibitory concentration of three quinones and two reactive oxygen species (ROS) for Dap-S and Dap-R paired strains of pair A and pair D. No growth difference between Dap-R and Dap-S paired strains in presence of any of three tested quinones (menadione, benzoquinone, ortho-benzoquinone) or two reactive oxygen species (hydrogen peroxide, paraquat) (two-tailed paired Student’s t-test except for pair A/orthobenzoquinone, hydrogen peroxide, paraquat (Mann-Whitney test), p>0.05 for all pairs). (D) *In vitro* survival of Dap-S and Dap-R paired strain of two pairs exposed to ortho-benzoquinone (16 mg/l) or to hydrogen peroxide (4 mM) for 1 hour, as determined by CFU counts. Two-tailed paired Student’s t-test, ortho-benzoquinone: pair A, p=0.1665; pair D, p=0.0144; hydrogen peroxide: pair A, p=0.0127; pair D, p=0.1947. Dap-S, daptomycin susceptible; Dap-R, Daptomycin resistant; MIC, minimum inhibitory concentration.

The mechanism by which phenoloxidase kills bacteria is not entirely clear; in addition to melanin, this reaction produces reactive oxygen and cytotoxic quinones as intermediates [30,31,53]. Phenoloxidases are thought to generate a direct antimicrobial activity through these toxic products [53]. We thus investigated whether the observed differential sensitivity to melanization could be attributed to differential susceptibility to either reactive oxygen species or quinones. Because the quinones generated during the melanization reaction (e.g. dopaquinone) are highly labile and thus not commercially-available, we investigated this question by assaying sensitivity *in vitro* to menadione, para-benzoquinone, and ortho-benzoquinone and to two reactive oxygen species (hydrogen peroxide, paraquat). We found no difference in minimum inhibitory concentration of any of these compounds (**Fig 7C**). Ortho-quinones, those produced by the melanization reaction, are normally considered the most reactive, so we further tested bacterial survival to ortho-benzoquinone. However, different pairs exhibited different changes in sensitivity to killing by ortho-benzoquinone, indicating that sensitivity to quinones is unlikely to represent the common mechanism driving loss of virulence in Dap-R strains (**Fig 7D**, Figure **S7B**). Collectively, our data demonstrate that the loss of virulence in daptomycin resistant strains reflects increased sensitivity to prophenoloxidase-dependent melanization *in vivo*.

## Discussion

When subjected to antimicrobial selective pressure, *S. aureus* acquires resistance which can affect its pathogenicity [19,54]. Using a *Drosophila* systemic infection model and a panel of paired clinical strains collected before and after daptomycin resistance acquisition, the largest so far reported, we show that acquisition of daptomycin resistance, most frequently through *mprF* mutation, is associated with attenuated pathogenicity that is mostly explained by greater susceptibility to *Drosophila* phenoloxidase activity. We also show that this attenuated virulence is independent of host detection via the Toll pathway, and that the virulence difference between daptomycin resistant and susceptible strains is not a consequence of interference with host defence peptide expression. Finally, we show that this acquired susceptibility to phenoloxidase activity is not a consequence of a general sensitivity to quinones but does require direct contact between *S. aureus* and phenoloxidase.

Most of our daptomycin resistant isolates differed from their daptomycin sensitive parental strains by the presence of a single mutation of unknown functional impact in *mprF*. Mutations that confer daptomycin resistance have been assumed to lead to *mprF* gain-of-function, resulting in increased abundance of lysyl-phosphatidylglycerol in the outer leaflet of the membrane [55–57]. This change would be expected to result in increased membrane charge, which might help repel cationic AMPs from the membrane, modulate interaction of peptides with the membrane, or attenuate membrane perturbation by daptomycin and potentially other antimicrobial peptides [10,58,59]. MprF is also assumed to also influence cell wall metabolism, membrane fluidity, membrane proteome, preformed toxin release, or general fitness [34,57,60]. However, consistent with other reports, we found no change in surface charge or in a range of other bacterial surface properties in our Dap-R strains [41,42,61,62]. The reasons for this apparent difference are poorly understood; they may be related to some unexamined difference in genetic background or subtleties of the precise mutations identified. Resistance to daptomycin may also result from an effect of MprF enzymatic activities other than those known (synthesis and transport of lysyl-phosphatidylglycerol) [63]. Finally, they may reflect some fundamental misunderstanding of the interaction between daptomycin and Mprf itself.

Despite this gap in our knowledge, what is clear is that *mprF*-related daptomycin resistance exhibits phenotypic crosstalk with staphylococcal pathogenic potential. Recently, it has been shown that blockade of the MprF flippase activity sensitized MRSA to host AMPs and daptomycin, which interfered with pathogenicity in humans [64]. Here, we found that *mprF* mutations, while conferring daptomycin resistance, also caused a decrease in virulence by increasing sensitivity to *Drosophila* host defence. The so-called ‘see-saw effect’ refers to the observation that increased resistance to daptomycin is associated with reduced resistance to β-lactam antibiotics [39]; our findings show a similar ‘see-saw’ between daptomycin resistance and pathogenic potential.

When we initiated this work, we first wished to assess whether daptomycin resistance could compromise fly AMP action, in line with previous reports of cross-resistance between daptomycin and cationic AMP such as human LL-37 or hNP-1 [26,42]. Neither the strong daptomycin resistant-associated attenuation of virulence nor the Toll signalling pathway-independent effect support the concept of cross-resistance between daptomycin and host AMPs (**Fig 1A**, **2A, S2 Fig**). *Drosophila* host defence against *S. aureus* is dependent on the activities of many antimicrobial effectors, including antimicrobial peptides, Bomanins (a family of peptides without demonstrated antimicrobial activity), and phenoloxidase [29,52,65]. Despite this wealth of effectors, our data clearly indicate that daptomycin resistance specifically increases susceptibility to killing via phenoloxidase. Phenoloxidase is the central activity in the melanisation reaction, one of the most immediate immune responses in the fly; it catalyses the oxidation of phenols (e.g., tyrosine) to quinones that spontaneously polymerize to form melanin [31]. While much is known on PPO (e.g., location of PPO storage inside the cell, PPO 3D structure is [53,66,67], the cascade by which PPO is activated into phenoloxidase [29], the steps of the melanisation reaction [31]), little is known about how phenoloxidase kills bacteria. One prominent view is that phenoloxidase activity generates highly-reactive quinones, and these quinones are the relevant antimicrobial activity [68]. Our daptomycin-resistant isolates do not generally exhibit increased sensitivity to quinones or ROS *in vitro*, but we were unable to directly test sensitivity to the specific quinones produced *in vivo* and thus cannot exclude increased sensitivity to these compounds in the host context.

Although we are unable to define the precise killing mechanisms of phenoloxidase, our observations have implications regarding how bacteria evade phenoloxidase-mediated killing. MprF has a well-defined role in modulating the staphylococcal cell surface. Here, we show that antistaphylococcal phenoloxidase activity is altered by the changes associated with MprF-related daptomycin resistance. Our data indicate that the cell-surface changes made by daptomycin-associated *mprF* alleles result in dramatically increased sensitivity to the bactericidal activity of phenoloxidase. Jearaphunt *et al* have demonstrated that agglutination accompanies, and may be the primary mechanism of, phenoloxidase-mediated killing [69]. It is possible that the effect of daptomycin resistance-associated *mprF* alleles is either to enable the association between circulating phenoloxidase and the bacterial surface or to enable the interaction between individual bacterial cells.

In a broader sense, our series of daptomycin resistant clinical paired isolates exhibits a strong and nearly constant trend of daptomycin resistance-associated virulence attenuation *in vivo*, consistent with previous observations in mouse, galleria, and zebrafish infection models [26,35]. The fact that daptomycin resistance is associated with reduced virulence in hosts with different immune systems, some of which lack antimicrobial phenoloxidase (e.g., zebrafish, mouse), is intriguing. One possibility is that this general effect is associated with a general difference in tendency to agglutinate. Another possibility is that the cell-surface modifications present in *mprF*-associated daptomycin resistance result in increased affinity for many immune effectors. Identification of the cell-surface properties responsible for this differential immune susceptibility is potentially a new staphylococcal Achilles heel.

## Methods

### Bacterial strains and culture conditions

*S. aureus* strains used in this study are listed in **S1 Table**. Strains were grown, unless specified, in 3 ml tryptic soy broth (TSB) in 30 ml tubes for 16 to 18 hours to stationary phase at 37°C with shaking (180 rpm). Where required, culture medium was supplemented with erythromycin (10 mg/ml). Enumeration of staphylococcal CFU was performed by plating 10-fold serial dilutions of samples onto tryptic soy agar (TSA) plates after an 18-hr incubation period at 37°C. Where required, TSA used for enumerations was supplemented with daptomycin at appropriate concentration (see competition assay) and CaCl_2_ was added to a final concentration of 1.25 mM.

### Bacterial growth assay

Stationary-phase bacteria were diluted 1000-fold in fresh broth into a 96-well flat-bottomed plate. The plate was incubated lid-covered into a TECAN Infinite 200 PRO microplate reader (Tecan Group Ltd., Switzerland). Bacteria were grown at 37°C with shaking (700 rpm), and bacterial growth was monitored using measurements of optical density at 600 nm (OD_600 nm_) every 15 min for a total of 18 h. Bacterial growth was successively assayed in TSB, cation-adjusted Muller-Hinton Broth (MHB), Dulbecco’s Modified Eagle Medium (DMEM) supplemented with 1% amino acids except for pair a strains (5% amino acids) and RPMI 1640.

### Genome sequencing and genome analysis

Whole genome sequencing (WGS) was performed for all strains and carried out at the GenePII Platform (Hospices Civils de Lyon, Lyon, France). DNA was extracted with the Maxwell RSC System (Promega, Madison, WI, USA) using the Maxwell RSC Blood DNA kit (Promega). Libraries were prepared using the DNA Prep library kit (Illumina, San Diego, CA, USA) and sequencing was done using a NextSeq 550 sequencer (Illumina) to obtain paired-end reads of 150 bp, respectively. Raw reads were trimmed for adapters and index removal and trimmed reads were used to produce assemblies with SPAdes v 3.14 [70]. Multi Locus Sequence Typing (MLST) was performed using assemblies (https://github.com/tseemann/mlst) and PubMLST (https://pubmlst.org/). Presence of point mutations associated to daptomycin was analysed using Point Finder and an inhouse curated database. Mutations were further corroborated by single nucleotide polymorphism (SNP) detection performed for each pair of clinical isolates collected before and after daptomycin resistance. Briefly, for each pair of clinical isolates, the genome of the daptomycin sensitive strain was annotated with Prokka v1.14.5 (https://github.com/tseemann/prokka) and used as the reference genome for the alignment of both trimmed reads from both daptomycin sensitive and resistant paired isolates. Core SNP detection was performed using Snippy v4.6.0 (https://github.com/tseemann/snippy). The sequence data for this study have been deposited in the European Nucleotide Archive (ENA) at EMBL-EBI under accession number PRJEB72842 and PRJEB54647 (for sample ERS12426074) (https://www.ebi.ac.uk/ena/browser/view/PRJEB72842 and https://www.ebi.ac.uk/ena/browser/view/PRJEB54647), and are listed in **S1 Table**.

### Determination of MICs

Two-fold dilutions of either antibiotic, quinone compounds, paraquat or hydrogen peroxide were prepared in cation-adjusted MHB in 96-well flat-bottomed plate. Stationary-phase *S. aureus* grown in TSB were diluted 1000-fold in MHB and inoculated into wells to a final concentration of 5 x 10^5^ CFU/ml bacteria. For daptomycin MICs determination, MHB was supplemented with CaCl_2_ to a final concentration of 1.25 mM. When oxacillin MICs were determined, MHB was supplemented with 2% NaCl. Determination of hydrogen peroxide, menadione, benzoquinone or ortho-benzoquinone MICs were performed with freshly prepared solutions. The 96-well was incubated statically at 37°C for 17 h. The MIC was determined as the lowest concentration of antibiotic with no visible growth. For purpose of clarity, we named daptomycin-resistant strains with daptomycin MIC >1 μg/mL even though the official term is daptomycin-non susceptible [71].

### Determination of bacterial survival to ortho-benzoquinone and hydrogen peroxide

Assays were based on Ha et al [72]. Briefly, stationary-phase bacterial cells were washed twice with PBS and adjusted to 10^6^ CFU/ml. 10 μl of the bacterial suspension (10^4^ CFU) was added to wells in a sterile 96-well flat-bottomed plate, and 90 μl of freshly diluted ortho-benzoquinone (16 mg/L in PBS), freshly diluted hydrogen peroxide (4 mM/L in PBS) or PBS (for 0 h time point) was added. The 96-well plate was incubated for 1 h at 37 °C in the dark without shaking. Bacterial survival to ortho-benzoquinone or to hydrogen peroxide was determined by enumerating CFU on TSA plates or on Columbia blood agar plates, respectively, after subjecting the suspensions to 10-fold serial dilutions. Results were expressed as a percentage of the number of bacteria in the initial inoculum.

### Determination of the hemolytic activity

Haemolysis of human erythrocytes was used to measure of haemolytic activity of *S. aureus* hemolysins, as described previously [73]. Ethical approval for drawing and processing human blood was obtained from the Regional Ethics Committee of Imperial College healthcare tissue bank (Imperial College London) and the Imperial NHS Trust Tissue Bank (REC Wales approval no. 12/WA/0196 and ICHTB HTA license no. 12275). Written informed consent was obtained from the donors prior to taking samples. Aliquots (300 ml) from bacterial cultures in TSB were sampled at various time points (0h, 2h, 3h, 4h, 5h, 6h). Bacterial cells were removed by centrifugation (17,000 x g, 10 min), and the supernatant were subjected to serial 2-fold dilutions in 96-well microplates in fresh TSB. Aliquots from each dilution (100 ml) were mixed with an equal volume of 5% human blood suspension in PBS. The 96-well plate was incubated statically at 37 °C for 1 h. The 96-well plates were then centrifuged (500 x g, 5 min) to remove the intact blood cells, and supernatants (100 ml) were transferred into the wells of a new 96-well flat-bottomed plate. The intensity of erythrocyte cell lysis was determined by measuring absorbance at 570 nm (A570) using an iMark^TM^ microplate reader (Bio-Rad) [72,73]. Blood incubated in TSB only served as a negative control, while TSB containing 0.1% Triton X-100 served as a positive control and was considered to represent 100% lysis. Percent haemolysis for each sample and time point was calculated relative to positive control. All values used in calculations were related to the A570 reading of undiluted supernatants from the bacterial cultures.

### Determination of *S. aureus* exoproteome

The content of secreted proteins was evaluated as described previously [74]. Bacteria were removed from 4-h spent culture medium by centrifugation (4,000 x g; 15 min). 30 ml of 40% trichloroacetic acid in acetone was added to an aliquot (10ml) of supernatant, and the tubes were incubated at -20°C overnight. Protein pellets were washed twice with acetone, concentrated 10-fold by centrifugation (4,000 x g; 15 min), and dried. The pellets were resuspended in sample electrophoresis treatment buffer (6% SDS, 15% glycerol, 3% b-mercaptoethanol, 0.02% bromophenol blue, 0.075 M Tris-HCL, pH 7). The samples were heated at 95°C for 10 min, centrifuged (17,000 x g, 1 min) and run on 10% SDS-polyacrylamide gels. After being stained for protein with Coomassie blue, the gels were washed and distained before being imaged on a Biorad Gel doc EZ imager with Image Lab 6.1 software (Bio-Rad).

### *Drosophila* stock and generation

Flies were maintained on a standard diet composed of 10% yeast (w/v), 8% fructose (w/v), 2% polenta (w/v), 0.8% w/v agar (w/v), 0.075% methylparaben (w/v) and 0.0825% propionic acid (v/v), as described elsewhere [75]. The fly lines used in this study are described in **S2 Table**. For crosses, flies were sheltered at 25°C, apart from flies expressing Toll^10b^ who were housed at room temperature until eclosion. After hatching, male flies were collected and kept at 25°C on fly medium in groups of 20 per vial. 5–8-day old male flies were used for all experiments. Unless specified otherwise, W^1118^ flies (hereafter wild type flies) were used for experiments.

### Drosophila injections

Fly injections were carried out as previously described [75]. Briefly, injections were performed with a Picospritzer III instrument (Parker, New Hampshire, US) and microinjection needles produced from borosilicate glass capillaries. CO_2_ anesthetized flies were injected in the lateral anterior of the abdomen with 50 nl of a standardized bacterial suspension in sterile PBS. Control flies included flies injected with sterile PBS and uninjected flies. After inoculation, flies were housed at 29 °C on standard diet.

For bacterial injection, bacterial suspensions were prepared as follows: bacteria grown to stationary-phase (15 h) were harvested by mild centrifugation (4,000 x g, 5 min) and cells were resuspended in PBS. The bacterial suspensions were adjusted to OD_600_ 0.1 in sterile PBS and further diluted 1:50 before fly injection, except for pair c (dilution 1:10). This resulted in an approximate average inoculum of 50 to 80 CFU in the fly body (hereafter standard inoculum). Infectious dose of every infection assay was checked by culturing serially diluted bacterial suspensions and two individual flies homogenized instantly after injection in 100 µl of sterile PBS and plated onto TSA plate.

For competition assays, flies were injected with a mixture of the paired strains at a ratio of 1:1. This mixture comprised a total amount of bacteria corresponding to the standard inoculum, and half of that inoculum was daptomycin susceptible strain while the other half was daptomycin resistant strain. Control flies included, in addition to the standard controls (PBS-injected, uninjected flies), flies injected with either individual strains (daptomycin susceptible strain or resistant strain) at standard inoculum or individual strain at 2-fold diluted standard inoculum (half standard inoculum). These controls were designed to compare the outcomes of monomicrobial and mixed infections with either the same total amount of bacteria (all cells cumulated) or with the same amount of each individual microorganism.

### Fly survival assays

For each experiment, a total of 20 adult flies per condition were injected and survival was regularly assessed for 6 days by scoring dead flies. At least two independent experiments were carried out.

### Drosophila gene expression -RT-qPCR

Samples were collected 3 h and 6 h after infection. Each sample comprised three flies that were homogenized in 100 µl TRI reagent (Sigma). Samples were stored at -20°C before nucleic acid extraction. RNA was isolated following the TRI reagent manufacturer’s protocol. Samples were subjected to a chloroform extraction and precipitation in isopropanol. The resultant pellet was washed with 70% ethanol and then subjected to RNAse-free DNAse treatment before DNAse was inactivated by addition of EDTA and heat treatment at 70°C. Aliquots (10 ml) were treated with RevertAid M-MuLV reverse transcriptase and random hexamers (Thermo Fisher Scientific) to synthetise complementary DNA (cDNA). Then, 10 µl of each cDNA sample (i.e., half volume of each sample) was pooled to generate 8 standards by 5-fold serial dilutions which were further used to generate a standard curve for each gene. Aliquot of every cDNA sample (10 ml) was 40-fold diluted before being used for gene quantification by qRT-PCR, using Sensimix with qPCR SyGreen 2x qPCR mix (PCR Biosystems) on a Corbett Rotor-Gene 6000. The cycling conditions were as follows: hold at 95°C for 10 min, then 45 cycles of 95°C for 15 s, 57°C for 30 s, and 72°C for 30 s, followed by a melting curve. Gene expression was determined from the standard curve generated during each run, normalised to the expression of housekeeping gene *RpL4*. Samples from PBS-injected and microbes-injected were then divided by the mean value of their uninjected controls to generate expression values relative to uninfected flies. Primers used are listed in **S3 Table**. Assay was performed twice with at least 3 biological replicates per condition per experiment.

### Bacterial quantification in the fly

At least 16 flies were injected with *S. aureus* daptomycin susceptible strain or its resistant paired strain. From these, eight flies were collected for 0 h quantifications while the remaining flies were housed at 29°C for 18 to 20 h (depending on the survival curve aspect of every pair) before homogenisation and bacterial quantification. In any case, individual flies were homogenized in 100 µl of sterile PBS. The entire homogenates (100 µl) were plated undiluted onto TSA for flies collected at 0 h, while samples from flies collected at 18-20 h were serially diluted and plated onto TSA plates. Plates were incubated for 16-18 h at 37°C. The number of colonies were counted, and calculation was processed to determine the number of CFUs present in each fly. All quantifications were performed twice.

For competition assay, flies were processed following this protocol. Briefly, the homogenates of flies infected with the mix (daptomycin susceptible and resistant paired strains) were plated onto TSA supplemented with CaCl_2_ to a final concentration of 1.25 mM and daptomycin (hereafter Dap-TSA) to final concentration of either 0.5 mg/L (pair a strains) or 1 mg/L (pair d strains). Following plate incubation and colony counting, the number of daptomycin resistant strain CFUs present in each fly were calculated from the number of colonies counted on the Dap-TSA plates, while the number of daptomycin susceptible strain CFUs were determined from the difference between the number of colonies on TSA (Daptomycin susceptible and resistant strain CFUs) and on Dap-TSA plates (daptomycin resistant strain CFUs). Because the homogenates (100 µl) of flies collected for bacterial quantification at 0 h were entirely spread onto plates, an extra 8 flies were injected with the mix (daptomycin susceptible and daptomycin paired strains) for quantifications at that time point, homogenized in PBS and the entire samples were plated on Dap-TSA before an overnight incubation at 37°C and colony counting.

### Determination of membrane fluidity

The fluidity of cell membrane was assayed by Laurdan generalized polarization. 500 ml stationary-phase culture (17 h) were incubated in the dark at room temperature for 5 min with Laurdan at a final concentration of 100 mM. Cells were then washed three time in PBS and 100 ml of the bacterial suspension were transferred into the wells of a black-walled 96-well plate. Laurdan fluorescence intensities were measured with a Tecan Infinite 200 Pro multiwell plate reader using an excitation wavelength of 330 nm, and emission wavelengths of 460 nm and 500 nm. Generalised polarisation was determined using the formula GP = (I460 – I500)/(I460+I500), where I460 and I500 are the emission intensity at 460 and 500 nm, respectively [76]. Assays were repeated at least 3 times. A higher GP value indicated a relatively more rigid membrane, while a lower GP value indicated a more fluid membrane.

### Determination of peptidoglycan production

The cell wall thickness was assessed using HCC-amino-d-alanine (HADA), a fluorescent d-amino acid analogue that is incorporated into peptidoglycan during its synthesis. Bacteria were seeded into 30 ml universal tubes containing 3 ml fresh TSB and HADA at a final concentration of 25 mM. Tubes were incubated in the dark for 16 hours to stationary phase at 37°C with shaking (180 rpm). The samples were washed three times in PBS. 200 μL of the bacterial suspension were transferred into the wells of a black-walled 96-well plate. HADA fluorescence intensities were measured using a Tecan Infinite 200 Pro multiwell plate reader using an excitation wavelength of 405 nm and emission wavelengths of 450 nm. Measurements generated values expressed as relative fluorescence units (RFU) correlate the amount of peptidoglycan produced, and indirectly, of the cell wall thickness during growth.

### Determination of surface charge

The charge of bacterial surface was measured using fluorescein isothiocyanate FITC-labelled Poly-L-Lysine (FITC-PLL) [77]. One ml of bacteria grown to stationary-phase overnight were washed and then incubated with FITC-PLL (final concentration 80 μg/mL) at room temperature for 10 min in the dark. The samples were washed three times in PBS, the cells were resuspended and 200 μL of the bacterial suspension were transferred into the wells of a black-walled 96-well plate. The fluorescence was measured using a TECAN Infinite PRO 200 plate reader using an excitation wavelength of 485 nm and emission wavelengths of 525 nm. Higher the signal, expressed as RFU, reflected an increased FITC-PLL binding to the bacterial surface, which indicated a more negatively cell surface charge. Controls (PBS, wild type USA 300 LAC*, USA 300 LAC JE2, USA 300 LAC JE2 *mprF*::Tn, USA 300 LAC* DdltD) were included in the series.

### Cell surface carotenoid extraction and quantification

Amounts of cell surface carotenoid content (including staphyloxanthin) were determined as previously described [72]. Cells were pelleted from aliquots (1 ml) of bacteria grown to stationary-phase overnight by centrifugation at 17,000 × *g* for 2 min. Cells were resuspended in methanol and incubated for 30 min at 42°C. Then, 100 μl of the supernatant was transferred to wells of a 96-well flat-bottomed plate. The absorbance was measured at 450 nm using a biorad iMark microplate reader to estimate the amount of extracted staphyloxanthin. Control strains (wild type USA 300 LAC JE2, USA 300 LAC JE2 *crtN*::Tn, USA 300 LAC JE2 *crtM*::Tn, SH1000) were included in the series.

### Statistical analysis

Experiments were performed on at least three independent occasions, unless specified otherwise. Data were analysed and figures were drafted in GraphPad Prism software version 9.5.1 (GraphPad Software Inc, USA). Error bars, where shown, represent the standard deviation of the mean. Boxplots represent median, percentile 25 and 75 and whiskers the minimum and maximum values. Survival curves were plotted using the Kaplan-Meier method. For single comparisons, a two-tailed Student’s t-test or Mann-Whitney test was used to analyse the data. For multiple comparisons, a one-way analysis of variance (ANOVA) or a Kruskal-Wallis test was performed. Survival data were analysed using the log-rank test. When appropriate, appropriate post hoc analysis (Tukey’s, Sidak’s, Dunn’s) was performed to identify specific differences between conditions. A *P* value of <0.05 was considered to reflect significance.

## Acknowledgements

Laurent Boyer (Inserm U1065) is warmly thanked for helpful advices in preliminary work.

## Author contribution

**Conceptualization**: Brigitte Lamy, Frédéric Laurent, Andrew M Edwards, Marc S Dionne

**Data curation**: Brigitte Lamy, Camille Kolenda

**Formal analysis**: Brigitte Lamy

**Funding acquisition**: Andrew M Edwards, Marc S Dionne, Frédéric Laurent, Brigitte Lamy

**Investigation**: Brigitte Lamy, Carolina Simoes Da Silva, Ashima Wadhawan, Camille Kolenda, Patricia Martins Simoes

**Methodology**: Andrew M Edwards, Marc S Dionne, Elisabeth Ledger

**Supervision**: Andrew M Edwards, Marc S Dionne

**Visualisation**: Brigitte Lamy

**Writing – original draft**: Brigitte Lamy, Marc S Dionne, Andrew M Edwards

**Writing – review & editing**: Brigitte Lamy, Andrew M Edwards, Marc S Dionne, Frédéric Laurent, Camille Kolenda, Ashima Wadhawan, Elizabeth Ledger, Carolina Simoes Da Silva, Patricia Martins Simoes

## Supporting information

**S1 Fig. Paired strains *in vitro* growth over time.** Growth in (A) tryptic soy broth; (B) Muller Hinton broth; (C) Dulbecco’s Modified Eagle Medium (DMEM) supplemented with amino acids; (D) RPMI 1640. In all cases, at least 3 replicates in duplicate or triplicate.

**S2 Fig. Cell surface charge correlates with neither daptomycin resistance nor virulence in the fly.** Cell surface charge of overnight TSB grown cells as determined by binding of highly positively charged fluorescein isothiocyanate-poly-L-lysine (FITC-PLL). Data from at least three independent experiments. Dashed lines link plots from same experiment. Data were analysed by a two-tailed paired Student’s t-test. Difference in cell surface charge between paired strains is evidenced for none pair (p>0.05, all pairs) but pair j (p=0.0295). TSB: Trypic soy broth; RFU: relative fluorescence units; Dap-S, daptomycin susceptible; Dap-R, daptomycin resistant.

**S3 Fig. Fluidity of cell membrane correlates with neither daptomycin resistance nor virulence in the fly.** Membrane fluidity was determined using the fluorescent Laurdan dye to generate Generalised Polarisation. Data from at least three independent experiments. Dashed lines link plots from same experiment. Data were analysed by a two-tailed unpaired Student’s t-test. Difference in membrane fluidity between paired strains is evidenced in none pair (p>0.05 for all pairs). RFU: relative fluorescence units; Dap-S, daptomycin susceptible; Dap-R, daptomycin resistant.

**S4 Fig. Cell wall thickness correlate with neither daptomycin resistance nor virulence in the fly.** Cell wall thickness was determined by the intake of a fluorescent peptidoglycan precursor (HADA) in cell wall of overnight grown cells in TSB with HADA. Data from at least three independent experiments are shown. Dashed lines link plots from same experiment. Data were analysed by a two-tailed unpaired Student’s t-test. No difference in cell wall thickness is evidenced between paired strains for pairs a-e, h,i, k (p>0.05) while a difference is observed for pair g (p=0.0330). TSB: Trypic soy broth; RFU: relative fluorescence units; Dap-S, daptomycin susceptible; Dap-R, Daptomycin resistant.

**S5 Fig. Cell surface carotenoid content correlates with neither daptomycin resistance nor virulence in the fly.** Data from three independent experiments. Dashed lines links data from same experiment. Data were analysed by a two-tailed unpaired Student’s t-test. Difference in cell surface carotenoid content between paired strains is evidenced for pair a (p=0.0267) and pair h (p=0.0341), for all other pairs p>0.05. RFU: relative fluorescence units; Dap-S, daptomycin susceptible; Dap-R, daptomycin resistant.

**S6 Fig. Bomanins do not affect staphylococcal virulence depending on daptomycin resistance.** Survival of flies infected with either Dap-S strain or Dap-R paired strain. Flies from two fly lineages were infected: wild type (*W^1118^*) and bomanin-deficient (*Bom^Δ55C^*, IM2) flies. Data presented for 2 pairs, pair a and pair d. Survival curves correspond to one assay representative of at least two independent experiments. Data were analysed using the log-rang test: pair a, p<0.0001 (*W^1118^*) and p<0.0001 (*Bom^Δ55C^*); pair d, p=0.0005 (*W^1118^*) and p=0.0125 (*Bom^Δ55C^*). Dap-S, daptomycin susceptible; Dap-R, Daptomycin resistant.

**S7 Fig. Daptomycin resistant staphylococcal associated virulence is affected by prophenoloxidase but cells are not more susceptible to quinones.** (A) Survival of wild type (*W^1118^*) and *W^1118^;; PPO1Δ, PPO2Δ* mutants flies infected with either Dap-S strain or Dap-R paired strain. Lifespan analysis shows a consistent reduced difference in mortality of *PPO1Δ, PPO2Δ* double mutant flies between animals infected with Dap-S and Dap-R strains compared to wild type flies. A notable exception concerns pairs j and k that exhibit limited difference in virulence of the Dap-S and Dap-R paired strains in the wild type flies. Survival curves representative of two independent experiments. *W^1118^*: log-rang test, Pairs b, c, f, and j, p≤0.0001, pair e, p=0.027, pair g, p=0.001, pair h, p=0.039, pair i: 0.15, pair k, p=0.22; PBS and UI, P>0.05. *W^1118^*: Pairs b, c, f, and j, p≤0.0001, pair e, p=0.027, pair g, p=0.001, pair h, p=0.039, pair i: 0.15, pair k, p=0.22; PBS and UI, P>0.05; *PPO1Δ, PPO2Δ:* Pairs b, c, f, g, and i, p≤0.0001, pair e, p=0.0002, pair h, p=0.0007, pair j, p=0.0018, pair k: 0.0565, PBS and UI, P>0.05. (B) *In vitro* Survival of Dap-S and Dap-R paired strain of pairs other than pairs a and d exposed to ortho-benzoquinone (16 mg/l) for 1 hour, as determined by CFU counts. Survival does not correlate with bacterial virulence in the fly (two-tailed paired Student’s t-test, p>0.05 for all pairs). Dap-S, daptomycin susceptible; Dap-R, Daptomycin resistant; CFU, colony forming unit.

**S1 Table. Bacterial strains used in this study.**

**S2 Table. Fly lines used in this study and their sources**

**S3 Table. List of RT-qPCR primers used for expression analysis of *Drosophila* antimicrobial effectors in this study**

## References

1. Hindy J-R, Quintero-Martinez JA, Lee AT, Scott CG, Gerberi DJ, Mahmood M, et al. Incidence Trends and Epidemiology of Staphylococcus aureus Bacteremia: A Systematic Review of Population-Based Studies. Cureus. 2022 [cited 2 Jan 2024]. doi:10.7759/cureus.25460

2. Bai AD, Lo CKL, Komorowski AS, Suresh M, Guo K, Garg A, et al. Staphylococcus aureus bacteraemia mortality: a systematic review and meta-analysis. Clinical Microbiology and Infection. 2022;28: 1076–1084. doi:10.1016/j.cmi.2022.03.015

3. Le Moing V, Alla F, Doco-Lecompte T, Delahaye F, Piroth L, Chirouze C, et al. Staphylococcus aureus Bloodstream Infection and Endocarditis - A Prospective Cohort Study. Greub G, editor. PLoS ONE. 2015;10: e0127385. doi:10.1371/journal.pone.0127385

4. Habib G, Lancellotti P, Antunes MJ, Bongiorni MG, Casalta J-P, Del Zotti F, et al. 2015 ESC Guidelines for the management of infective endocarditis: The Task Force for the Management of Infective Endocarditis of the European Society of Cardiology (ESC)Endorsed by: European Association for Cardio-Thoracic Surgery (EACTS), the European Association of Nuclear Medicine (EANM). Eur Heart J. 2015;36: 3075–3128. doi:10.1093/eurheartj/ehv319

5. Tabah A, Laupland KB. Update on Staphylococcus aureus bacteraemia. Curr Opin Crit Care. 2022;28: 495–504. doi:10.1097/MCC.0000000000000974

6. Delgado V, Ajmone Marsan N, De Waha S, Bonaros N, Brida M, Burri H, et al. 2023 ESC Guidelines for the management of endocarditis. European Heart Journal. 2023;44: 3948–4042. doi:10.1093/eurheartj/ehad193

7. Galar A, Weil AA, Dudzinski DM, Muñoz P, Siedner MJ. Methicillin-Resistant Staphylococcus aureus Prosthetic Valve Endocarditis: Pathophysiology, Epidemiology, Clinical Presentation, Diagnosis, and Management. Clin Microbiol Rev. 2019;32: e00041–18. doi:10.1128/CMR.00041-18

8. Holubar M, Meng L, Alegria W, Deresinski S. Bacteremia due to Methicillin-Resistant Staphylococcus aureus: An Update on New Therapeutic Approaches. Infect Dis Clin North Am. 2020;34: 849–861. doi:10.1016/j.idc.2020.04.003

9. Ji S, Jiang S, Wei X, Sun L, Wang H, Zhao F, et al. In-Host Evolution of Daptomycin Resistance and Heteroresistance in Methicillin-Resistant Staphylococcus aureus Strains From Three Endocarditis Patients. The Journal of Infectious Diseases. 2020;221: S243–S252. doi:10.1093/infdis/jiz571

10. Thitiananpakorn K, Aiba Y, Tan X-E, Watanabe S, Kiga K, Sato’o Y, et al. Association of mprF mutations with cross-resistance to daptomycin and vancomycin in methicillin-resistant Staphylococcus aureus (MRSA). Sci Rep. 2020;10: 16107. doi:10.1038/s41598-020-73108-x

11. Andersson DI, Hughes D. Antibiotic resistance and its cost: is it possible to reverse resistance? Nat Rev Microbiol. 2010;8: 260–271. doi:10.1038/nrmicro2319

12. Cameron DR, Howden BP, Peleg AY. The Interface Between Antibiotic Resistance and Virulence in Staphylococcus aureus and Its Impact Upon Clinical Outcomes. Clinical Infectious Diseases. 2011;53: 576–582. doi:10.1093/cid/cir473

13. Geisinger E, Isberg RR. Interplay Between Antibiotic Resistance and Virulence During Disease Promoted by Multidrug-Resistant Bacteria. The Journal of Infectious Diseases. 2017;215: S9– S17. doi:10.1093/infdis/jiw402

14. Linzner N, Fritsch VN, Busche T, Tung QN, Loi VV, Bernhardt J, et al. The plant-derived naphthoquinone lapachol causes an oxidative stress response in Staphylococcus aureus. Free Radical Biology and Medicine. 2020;158: 126–136. doi:10.1016/j.freeradbiomed.2020.07.025

15. Yamaguchi T, Ando R, Matsumoto T, Ishii Y, Tateda K. Association between cell growth and vancomycin resistance in clinical community-associated methicillin-resistant Staphylococcus aureus. IDR. 2019;Volume 12: 2379–2390. doi:10.2147/IDR.S209591

16. Cui L, Ma X, Sato K, Okuma K, Tenover FC, Mamizuka EM, et al. Cell Wall Thickening Is a Common Feature of Vancomycin Resistance in *Staphylococcus aureus*. J Clin Microbiol. 2003;41: 5–14. doi:10.1128/JCM.41.1.5-14.2003

17. Cui J, Zhang H, Mo Z, Yu M, Liang Z. Cell wall thickness and the molecular mechanism of heterogeneous vancomycin-intermediate *Staphylococcus aureus*. Lett Appl Microbiol. 2021;72: 604–609. doi:10.1111/lam.13456

18. Howden BP, Smith DJ, Mansell A, Johnson PDR, Ward PB, Stinear TP, et al. Different bacterial gene expression patterns and attenuated host immune responses are associated with the evolution of low-level vancomycin resistance during persistent methicillin-resistant Staphylococcus aureus bacteraemia. BMC Microbiol. 2008;8: 39. doi:10.1186/1471-2180-8-39

19. Cameron DR, Lin Y-H, Trouillet-Assant S, Tafani V, Kostoulias X, Mouhtouris E, et al. Vancomycin-intermediate Staphylococcus aureus isolates are attenuated for virulence when compared with susceptible progenitors. Clinical Microbiology and Infection. 2017;23: 767–773. doi:10.1016/j.cmi.2017.03.027

20. Novick RP, Geisinger E. Quorum Sensing in Staphylococci. Annu Rev Genet. 2008;42: 541–564. doi:10.1146/annurev.genet.42.110807.091640

21. Salemi R, Zega A, Aguglia E, Lo Verde F, Pigola G, Stefani S, et al. Balancing the Virulence and Antimicrobial Resistance in VISA DAP-R CA-MRSA Superbug. Antibiotics. 2022;11: 1159. doi:10.3390/antibiotics11091159

22. Taylor SD, Palmer M. The action mechanism of daptomycin. Bioorganic & Medicinal Chemistry. 2016;24: 6253–6268. doi:10.1016/j.bmc.2016.05.052

23. Gray DA, Wenzel M. More Than a Pore: A Current Perspective on the In Vivo Mode of Action of the Lipopeptide Antibiotic Daptomycin. Antibiotics. 2020;9: 17. doi:10.3390/antibiotics9010017

24. Mishra NN, Yang S-J, Chen L, Muller C, Saleh-Mghir A, Kuhn S, et al. Emergence of Daptomycin Resistance in Daptomycin-Naïve Rabbits with Methicillin-Resistant Staphylococcus aureus Prosthetic Joint Infection Is Associated with Resistance to Host Defense Cationic Peptides and mprF Polymorphisms. Becker K, editor. PLoS ONE. 2013;8: e71151. doi:10.1371/journal.pone.0071151

25. Sakoulas G, Guram K, Reyes K, Nizet V, Zervos M. Human Cathelicidin LL-37 Resistance and Increased Daptomycin MIC in Methicillin-Resistant Staphylococcus aureus Strain USA600 (ST45) Are Associated with Increased Mortality in a Hospital Setting. Diekema DJ, editor. J Clin Microbiol. 2014;52: 2172–2174. doi:10.1128/JCM.00189-14

26. Bhuiyan MS, Jiang J-H, Kostoulias X, Theegala R, Lieschke GJ, Peleg AY. The Resistance to Host Antimicrobial Peptides in Infections Caused by Daptomycin-Resistant Staphylococcus aureus. Antibiotics. 2021;10: 96. doi:10.3390/antibiotics10020096

27. Lemaitre B, Hoffmann J. The host defense of Drosophila melanogaster. Annu Rev Immunol. 2007;25: 697–743. doi:10.1146/annurev.immunol.25.022106.141615

28. Filipe SR, Tomasz A, Ligoxygakis P. Requirements of peptidoglycan structure that allow detection by the Drosophila Toll pathway. EMBO Rep. 2005;6: 327–333. doi:10.1038/sj.embor.7400371

29. Dudzic JP, Hanson MA, Iatsenko I, Kondo S, Lemaitre B. More Than Black or White: Melanization and Toll Share Regulatory Serine Proteases in Drosophila. Cell Reports. 2019;27: 1050–1061.e3. doi:10.1016/j.celrep.2019.03.101

30. Jiang J, Zhou Z, Dong Y, Cong C, Guan X, Wang B, et al. In vitro antibacterial analysis of phenoloxidase reaction products from the sea cucumber Apostichopus japonicus. Fish & Shellfish Immunology. 2014;39: 458–463. doi:10.1016/j.fsi.2014.06.002

31. Whitten M, Coates CJ. Re-evaluation of insect melanogenesis research: Views from the dark side. Pigment Cell Melanoma Res. 2017; 386–401. doi:10.1111/pcmr.12590

32. Yeaman MR, Yount NY. Mechanisms of antimicrobial peptide action and resistance. Pharmacol Rev. 2003;55: 27–55. doi:10.1124/pr.55.1.2

33. Huang HW. DAPTOMYCIN, its membrane-active mechanism vs. that of other antimicrobial peptides. Biochimica et Biophysica Acta (BBA) - Biomembranes. 2020;1862: 183395. doi:10.1016/j.bbamem.2020.183395

34. Li S, Yin Y, Chen H, Wang Q, Wang X, Wang H. Fitness Cost of Daptomycin-Resistant Staphylococcus aureus Obtained from in Vitro Daptomycin Selection Pressure. Front Microbiol. 2017;8: 2199. doi:10.3389/fmicb.2017.02199

35. Taglialegna A, Varela MC, Rosato RR, Rosato AE. VraSR and Virulence Trait Modulation during Daptomycin Resistance in Methicillin-Resistant *Staphylococcus aureus* Infection. Fey PD, editor. mSphere. 2019;4: e00557–18. doi:10.1128/mSphere.00557-18

36. Peleg AY, Miyakis S, Ward DV, Earl AM, Rubio A, Cameron DR, et al. Whole Genome Characterization of the Mechanisms of Daptomycin Resistance in Clinical and Laboratory Derived Isolates of Staphylococcus aureus. Ahmed N, editor. PLoS ONE. 2012;7: e28316. doi:10.1371/journal.pone.0028316

37. Yang S-J, Nast CC, Mishra NN, Yeaman MR, Fey PD, Bayer AS. Cell Wall Thickening Is Not a Universal Accompaniment of the Daptomycin Nonsusceptibility Phenotype in *Staphylococcus aureus* : Evidence for Multiple Resistance Mechanisms. Antimicrob Agents Chemother. 2010;54: 3079–3085. doi:10.1128/AAC.00122-10

38. Mehta S, Cuirolo AX, Plata KB, Riosa S, Silverman JA, Rubio A, et al. VraSR Two-Component Regulatory System Contributes to *mprF* -Mediated Decreased Susceptibility to Daptomycin in *In Vivo* -Selected Clinical Strains of Methicillin-Resistant Staphylococcus aureus. Antimicrob Agents Chemother. 2012;56: 92–102. doi:10.1128/AAC.00432-10

39. Renzoni A, Kelley WL, Rosato RR, Martinez MP, Roch M, Fatouraei M, et al. Molecular Bases Determining Daptomycin Resistance-Mediated Resensitization to β-Lactams (Seesaw Effect) in Methicillin-Resistant Staphylococcus aureus. Antimicrob Agents Chemother. 2017;61: e01634–16. doi:10.1128/AAC.01634-16

40. Sulaiman JE, Lam H. Novel Daptomycin Tolerance and Resistance Mutations in Methicillin-Resistant Staphylococcus aureus from Adaptive Laboratory Evolution. Bradford PA, editor. mSphere. 2021;6: e00692–21. doi:10.1128/mSphere.00692-21

41. Ernst CM, Peschel A. MprF-mediated daptomycin resistance. International Journal of Medical Microbiology. 2019;309: 359–363. doi:10.1016/j.ijmm.2019.05.010

42. Bayer AS, Mishra NN, Chen L, Kreiswirth BN, Rubio A, Yang S-J. Frequency and Distribution of Single-Nucleotide Polymorphisms within *mprF* in Methicillin-Resistant Staphylococcus aureus Clinical Isolates and Their Role in Cross-Resistance to Daptomycin and Host Defense Antimicrobial Peptides. Antimicrob Agents Chemother. 2015;59: 4930–4937. doi:10.1128/AAC.00970-15

43. Bellido JLM. Mechanisms of resistance to daptomycin in. Rev Esp Quimioter.

44. Ledger EVK, Sabnis A, Edwards AM. Polymyxin and lipopeptide antibiotics: membrane-targeting drugs of last resort: This article is part of the Bacterial Cell Envelopes collection. Microbiology. 2022;168. doi:10.1099/mic.0.001136

45. Jenul C, Horswill AR. Regulation of *Staphylococcus aureus* Virulence. Fischetti VA, Novick RP, Ferretti JJ, Portnoy DA, Braunstein M, Rood JI, editors. Microbiol Spectr. 2019;7: 7.2.29. doi:10.1128/microbiolspec.GPP3-0031-2018

46. Thomer L, Schneewind O, Missiakas D. Pathogenesis of *Staphylococcus aureus* Bloodstream Infections. Annu Rev Pathol Mech Dis. 2016;11: 343–364. doi:10.1146/annurev-pathol-012615-044351

47. Needham AJ, Kibart M, Crossley H, Ingham PW, Foster SJ. Drosophila melanogaster as a model host for Staphylococcus aureus infection. Microbiology (Reading). 2004;150: 2347–2355. doi:10.1099/mic.0.27116-0

48. Wu K, Conly J, Surette M, Sibley C, Elsayed S, Zhang K. Assessment of virulence diversity of methicillin-resistant Staphylococcus aureus strains with a Drosophila melanogaster infection model. BMC Microbiol. 2012;12: 274. doi:10.1186/1471-2180-12-274

49. Meng X, Khanuja BS, Ip YT. Toll receptor-mediated Drosophila immune response requires Dif, an NF-kappaB factor. Genes Dev. 1999;13: 792–797. doi:10.1101/gad.13.7.792

50. Clemmons AW, Lindsay SA, Wasserman SA. An Effector Peptide Family Required for Drosophila Toll-Mediated Immunity. Silverman N, editor. PLoS Pathog. 2015;11: e1004876. doi:10.1371/journal.ppat.1004876

51. Maxton-Kuchenmeister Maxton-Kü null, Handel K, Schmidt-Ott U, Roth S, JackleJäckle H. Toll homologue expression in the beetle tribolium suggests a different mode of dorsoventral patterning than in drosophila embryos. Mech Dev. 1999;83: 107–114. doi:10.1016/s0925-4773(99)00041-6

52. Dudzic JP, Kondo S, Ueda R, Bergman CM, Lemaitre B. Drosophila innate immunity: regional and functional specialization of prophenoloxidases. BMC Biol. 2015;13: 81. doi:10.1186/s12915-015-0193-6

53. Lu A, Zhang Q, Zhang J, Yang B, Wu K, Xie W, et al. Insect prophenoloxidase: the view beyond immunity. Front Physiol. 2014;5. doi:10.3389/fphys.2014.00252

54. Gao W, Cameron DR, Davies JK, Kostoulias X, Stepnell J, Tuck KL, et al. The RpoB H481Y Rifampicin Resistance Mutation and an Active Stringent Response Reduce Virulence and Increase Resistance to Innate Immune Responses in Staphylococcus aureus. The Journal of Infectious Diseases. 2013;207: 929–939. doi:10.1093/infdis/jis772

55. Rubio A, Moore J, Varoglu M, Conrad M, Chu M, Shaw W, et al. LC-MS/MS characterization of phospholipid content in daptomycin-susceptible and -resistant isolates of *Staphylococcus aureus* with mutations in *mprF*. Molecular Membrane Biology. 2012;29: 1–8. doi:10.3109/09687688.2011.640948

56. Mishra NN, Bayer AS. Correlation of Cell Membrane Lipid Profiles with Daptomycin Resistance in Methicillin-Resistant Staphylococcus aureus. Antimicrob Agents Chemother. 2013;57: 1082– 1085. doi:10.1128/AAC.02182-12

57. Nakamura M, Kawada H, Uchida H, Takagi Y, Obata S, Eda R, et al. Single nucleotide polymorphism leads to daptomycin resistance causing amino acid substitution—T345I in MprF of clinically isolated MRSA strains. Narayanasamy P, editor. PLoS ONE. 2021;16: e0245732. doi:10.1371/journal.pone.0245732

58. Yang S-J, Mishra NN, Rubio A, Bayer AS. Causal Role of Single Nucleotide Polymorphisms within the *mprF* Gene of Staphylococcus aureus in Daptomycin Resistance. Antimicrob Agents Chemother. 2013;57: 5658–5664. doi:10.1128/AAC.01184-13

59. Song D, Jiao H, Liu Z. Phospholipid translocation captured in a bifunctional membrane protein MprF. Nat Commun. 2021;12: 2927. doi:10.1038/s41467-021-23248-z

60. Zheng X, Marsman G, Lacey KA, Chapman JR, Goosmann C, Ueberheide BM, et al. The cell envelope of Staphylococcus aureus selectively controls the sorting of virulence factors. Nat Commun. 2021;12: 6193. doi:10.1038/s41467-021-26517-z

61. Ernst CM, Peschel A. Broad-spectrum antimicrobial peptide resistance by MprF-mediated aminoacylation and flipping of phospholipids. Mol Microbiol. 2011;80: 290–299. doi:10.1111/j.1365-2958.2011.07576.x

62. Bayer AS, Mishra NN, Cheung AL, Rubio A, Yang S-J. Dysregulation of *mprF* and *dltABCD* expression among daptomycin-non-susceptible MRSA clinical isolates. J Antimicrob Chemother. 2016;71: 2100–2104. doi:10.1093/jac/dkw142

63. Sievers S, M. Ernst C, Geiger T, Hecker M, Wolz C, Becher D, et al. Changing the phospholipid composition of *Staphylococcus aureus* causes distinct changes in membrane proteome and membrane-sensory regulators. Proteomics. 2010;10: 1685–1693. doi:10.1002/pmic.200900772

64. Slavetinsky CJ, Hauser JN, Gekeler C, Slavetinsky J, Geyer A, Kraus A, et al. Sensitizing Staphylococcus aureus to antibacterial agents by decoding and blocking the lipid flippase MprF. eLife. 2022;11: e66376. doi:10.7554/eLife.66376

65. Hanson MA, Dostálová A, Ceroni C, Poidevin M, Kondo S, Lemaitre B. Synergy and remarkable specificity of antimicrobial peptides in vivo using a systematic knockout approach. eLife. 2019;8: e44341. doi:10.7554/eLife.44341

66. Li Y, Wang Y, Jiang H, Deng J. Crystal structure of *Manduca sexta* prophenoloxidase provides insights into the mechanism of type 3 copper enzymes. Proc Natl Acad Sci USA. 2009;106: 17002–17006. doi:10.1073/pnas.0906095106

67. Hu Y, Wang Y, Deng J, Jiang H. The structure of a prophenoloxidase (PPO) from Anopheles gambiae provides new insights into the mechanism of PPO activation. BMC Biol. 2016;14: 2. doi:10.1186/s12915-015-0225-2

68. Binggeli O, Neyen C, Poidevin M, Lemaitre B. Prophenoloxidase Activation Is Required for Survival to Microbial Infections in Drosophila. Schneider DS, editor. PLoS Pathog. 2014;10: e1004067. doi:10.1371/journal.ppat.1004067

69. Jearaphunt M, Noonin C, Jiravanichpaisal P, Nakamura S, Tassanakajon A, Söderhäll I, et al. Caspase-1-like regulation of the proPO-system and role of ppA and caspase-1-like cleaved peptides from proPO in innate immunity. PLoS Pathog. 2014;10: e1004059. doi:10.1371/journal.ppat.1004059

70. Prjibelski A, Antipov D, Meleshko D, Lapidus A, Korobeynikov A. Using SPAdes De Novo Assembler. Current Protocols in Bioinformatics. 2020;70: e102. doi:10.1002/cpbi.102

71. Jiang J-H, Bhuiyan MS, Shen H-H, Cameron DR, Rupasinghe TWT, Wu C-M, et al. Antibiotic resistance and host immune evasion in *Staphylococcus aureus* mediated by a metabolic adaptation. Proc Natl Acad Sci USA. 2019;116: 3722–3727. doi:10.1073/pnas.1812066116

72. Ha KP, Clarke RS, Kim G-L, Brittan JL, Rowley JE, Mavridou DAI, et al. Staphylococcal DNA Repair Is Required for Infection. mBio. 2020;11: 1–18. doi:10.1128/mBio.02288-20

73. Pader V, James EH, Painter KL, Wigneshweraraj S, Edwards AM. The Agr Quorum-Sensing System Regulates Fibronectin Binding but Not Hemolysis in the Absence of a Functional Electron Transport Chain. Fang FC, editor. Infect Immun. 2014;82: 4337–4347. doi:10.1128/IAI.02254-14

74. Smith DS, Siggins MK, Gierula M, Pichon B, Turner CE, Lynskey NN, et al. Identification of commonly expressed exoproteins and proteolytic cleavage events by proteomic mining of clinically relevant UK isolates of Staphylococcus aureus. Microbial Genomics. 2016;2. doi:10.1099/mgen.0.000049

75. Wadhawan A, Simoes Da Silva CJ, Nunes catarina, Edwards AM, Dionne MS. E. faecalis acquires resistance to antimicrobials and insect immunity via common mechanisms. bioRxiv 20220817504265; doi: 10.1101/20220817504265.

76. Müller A, Wenzel M, Strahl H, Grein F, Saaki TNV, Kohl B, et al. Daptomycin inhibits cell envelope synthesis by interfering with fluid membrane microdomains. Proc Natl Acad Sci USA. 2016;113. doi:10.1073/pnas.1611173113

77. Jones T, Yeaman MR, Sakoulas G, Yang S-J, Proctor RA, Sahl H-G, et al. Failures in clinical treatment of Staphylococcus aureus Infection with daptomycin are associated with alterations in surface charge, membrane phospholipid asymmetry, and drug binding. Antimicrob Agents Chemother. 2008;52: 269–278. doi:10.1128/AAC.00719-07

